# Integration of genetic, transcriptomic, and clinical data provides insight into 16p11.2 and 22q11.2 CNV genes

**DOI:** 10.1101/2020.06.23.166181

**Authors:** Mikhail Vysotskiy, Xue Zhong, Tyne W. Miller-Fleming, Dan Zhou, Autism Working Group of the Psychiatric Genomics Consortium, Bipolar Disorder Working Group of the Psychiatric Genomics Consortium, Schizophrenia Working Group of the Psychiatric Genomics Consortium, Nancy J. Cox, Lauren A Weiss

## Abstract

Deletions and duplications of the multigenic 16p11.2 and 22q11.2 copy number variants (CNVs) are associated with brain-related disorders including schizophrenia, intellectual disability, obesity, bipolar disorder, and autism spectrum disorder (ASD). The contribution of individual CNV genes to each of these phenotypes is unknown, as is the contribution of CNV genes to subtler health impacts. Hypothesizing that DNA copy number acts via RNA expression, we attempted a novel *in silico* fine-mapping approach in non-carriers using both GWAS and biobank data. We first asked whether expression level of a CNV gene impacts risk for a known brain-related phenotype(s). Using transcriptomic imputation, we tested for association within GWAS for schizophrenia, IQ, BMI, bipolar disorder, and ASD. We found individual genes in 16p11.2 associated with schizophrenia, BMI, and IQ (*SPN*), using conditional analysis to identify *INO80E* as the driver of schizophrenia, and *SPN* and *INO80E* as drivers of BMI. Second, we used a biobank containing electronic health data to compare the medical phenome of CNV carriers to controls within 700,000 individuals to investigate a spectrum of health effects, identifying novel and previously observed traits. Third, we used genotypes for over 48,000 biobank individuals to perform phenome-wide association studies between imputed expressions of 16p11.2 and 22q11.2 genes and over 1,500 health traits, finding seventeen significant gene-trait pairs, including psychosis (*NPIPB11, SLX1B*) and mood disorders (*SCARF2*), and overall enrichment of mental traits. Our results demonstrate how integration of genetic and clinical data aids in understanding CNV gene function, and implicate pleiotropy and multigenicity in CNV biology.

## INTRODUCTION

Multi-gene copy number variants (CNVs), including a 600kb region at 16p11.2 and a 3Mb region at 22q11.2, are known causes of multiple brain-related disorders. The 16p11.2 CNV, originally identified as a risk factor for autism spectrum disorder (ASD), has also been associated with schizophrenia, bipolar disorder, intellectual disability, and obesity^1–5^. The 22q11.2 CNV, identified as the cause of DiGeorge (velocardiofacial) syndrome, is associated with schizophrenia, intellectual disability, obesity, bipolar disorder, and ASD, as well ^6–11^. The effects of these two CNVs can be further subdivided into the effects of deletions vs. duplications. Some disorders are shared among carriers of deletions and duplications of the same region and others show opposite associations. For instance, ASD and intellectual disability are observed in both deletion and duplication carriers in both 16p11.2 and 22q11.2 ^3–8, 12–14^. Other traits are specific to one direction of the copy number change: schizophrenia and bipolar disorder are observed in 16p11.2 duplication carriers, but not deletion carriers ^15^. A third category of 16p11.2 and 22q11.2-associated traits are “mirrored”. 16p11.2 deletion carriers show increased rates of obesity, while duplication carriers tend to be underweight. 22q11.2 duplication carriers show reduced rates of schizophrenia, as opposed to increased rates in deletion carriers ^1,16, 17^. The question of which specific genes drive which brain-related traits associated with 16p11.2 or 22q11.2 CNVs remains unanswered. Likewise, what else these genes might be doing that has been more difficult to detect in small numbers of identified CNV carriers, who are primarily children? Identifying the role of specific gene(s) in behavioral and medical traits will clarify the biological processes that go awry as a result of these CNV mutations and the mechanisms by which they do so. Knowledge of the genes and mechanisms involved would, in turn, provide opportunities to develop targeted treatments.

Three of the traditional ways to map CNV genes to disorders are identifying loss-of-function mutations in these genes, analyzing smaller subsets of the entire region, and finding mutations in animal models that are sufficient to recapitulate the phenotype. The loss-of-function mutation method was used to fine-map the 17p11.2 CNV, another CNV associated with behavioral and non-behavioral traits ^18, 19^. Most of the features of the deletion syndrome, including intellectual disability, are represented in individuals who carry a defective copy of the *RAI1* gene due to point mutation ^20^. Duplications of *Rai1* appear to explain body weight and behavior abnormalities in mouse models of 17p11.2 duplications ^21^. Another example is the Williams syndrome CNV at 7q11.23 ^22, 23^. The cardiac traits associated with this syndrome are present in individuals with only one functional copy of the *ELN* gene, but this gene does not explain the behavioral traits ^24, 25^. The second method, of finding a smaller “critical region” was used to fine-map the 17q21.31 CNV ^26, 27^. By comparing patients who had similar symptoms with overlapping cytogenetic profiles, the common breakpoints of the CNV region were refined to a region containing only six genes ^27^. Later, Koolen *et al* identified patients showing intellectual disability and facial dysmorphisms characteristic of this CNV with disruptive mutations in one of the six genes, *KANSL1* ^28^. The third method of recapitulating similar phenotypes in animal models was successful in identifying *TBX1* as a gene important for some of the physical traits involved with 22q11.2 deletions. Mice with heterozygous mutations in the *TBX1* gene show cardiac outflow tract anomalies, similar to human 22q11.2 deletion carriers ^29–31^. However, it is unclear that *TBX1* is sufficient to explain brain-related disorders in 22q11.2 carriers ^32, 33^.

The 16p11.2 and 22q11.2 CNVs have been resistant to these traditional approaches for fine-mapping of brain-related traits. To date, no highly penetrant point mutations in 16p11.2 or 22q11.2 genes have been shown to be sufficient for a brain-related disorder. The most recent schizophrenia GWAS from the Psychiatric Genomics Consortium discovered a common SNP association near the 16p11.2 region, however the specific genes underlying GWAS signals are often unknown^34^. No small subsets of 16p11.2 or 22q11.2 genes have been proven necessary and sufficient to cause a brain-related disorder. A subregion of 22q11.2 has been proposed to explain ASD associated with deletions ^35^. As this subset of 22q11.2 contains approximately 20 genes, it is likely that further fine-mapping within this subset is possible. At 16p11.2, a subset of five deleted genes was isolated in a family with a history of ASD ^36^. However, this mutation neither caused ASD in all deletion carriers, nor was responsible for ASD in some non-carrier family members. Non-human models for the 16p11.2 and 22q11.2 CNVs, as well as knockouts for individual genes are available in mouse, zebrafish, and fruit flies ^37–42^, but have not successfully mapped individual genes in these CNVs to brain-related traits ^29–31^. Different zebrafish studies of 16p11.2 homologs have implicated different genes as phenotype drivers, as well as showed that most were involved in nervous system development ^38, 39, 43^. The complex brain-related traits associated with these CNVs are unlikely to be fully captured in model organisms. Hallucinations, a common symptom of schizophrenia, can be identified only in humans. There may be other aspects of 16p11.2 and 22q11.2 CNV biology that are human-specific. For example, mice carrying 16p11.2 duplications are obese, while obesity is associated with deletions in humans ^44^. Given the insufficiency of previous approaches, new approaches for fine-mapping genes in these regions to brain-related traits are necessary.

The motivation behind our approach is that in 16p11.2 and 22q11.2 CNV carriers, variation in gene copy number is expected to lead to variation in RNA expression level (with possible downstream effects on protein product). Expression measurements in mouse or human cell lines carrying 16p11.2 and 22q11.2 deletions and duplications confirm that for nearly all genes, duplication carriers have increased expression of individual CNV genes compared to controls and deletion carriers have reduced expression compared to controls ^45–50^. As the breakpoints of these CNVs do not disrupt the coding regions of individual genes, we believe that the variation in expression of one or more of the genes is the cause of pathogenicity. While these CNVs significantly disrupt gene expression levels, most genes’ expression levels vary among the general population, sometimes by a factor of two or more, as studies such as the Genotype-Tissue Expression Consortium (GTEx) have shown ^51–54^. This variation can be, in part, attributed to common genetic polymorphisms (expression quantitative trait loci, eQTLs). If large expression deviation in duplication and deletion carriers is a risk factor for a disorder, we hypothesize that more modest expression variation in the same genes among non-carriers will be a modest risk factor for the same disorder or milder related traits. This idea is analogous to the observation that common polymorphisms of small effect that are associated with a common trait can overlap with Mendelian genes for a similar trait ^55–57^.

Here, we perform three *in silico* studies of the impact of predicted expression of individual 16p11.2 and 22q11.2 genes, in comparison with the diagnosed CNVs, on human traits (Figure 1). First, we identify genes that associated with brain-related disorders via expression variation. Recent tools have leveraged the heritability of gene expression, allowing us to “impute” gene expression for genotyped individuals using eQTLs ^58, 59^. We perform association testing between imputed expression and five brain-related traits common to the 16p11.2 and 22q11.2 CNVs for which large amounts of genetic data have been amassed: schizophrenia, IQ, BMI, bipolar disorder, and ASD ^60–64^. We find at least one 16p11.2 gene associated with schizophrenia, IQ, and BMI. Second, we use a biobank containing electronic health records (EHRs) for over 3 million individuals to determine the medical traits in CNV carriers detected in our EHR system, confirming canonical CNV features and discovering novel over-represented traits ^65^. We also probe the consequences of expression variation of individual 16p11.2 and 22q11.2 genes on the medical phenome, by imputing gene expression in the >48,000 genotyped individuals in the same health system and performing a phenome-wide association test across all available traits. We find that mental disorders are over-represented among top gene-trait association pairs, and we highlight genes associated with the traits over-represented in CNV carriers. Taken together, our work provides a comprehensive catalog of associations of individual CNV genes to traits across the phenome.

**Figure 1:**
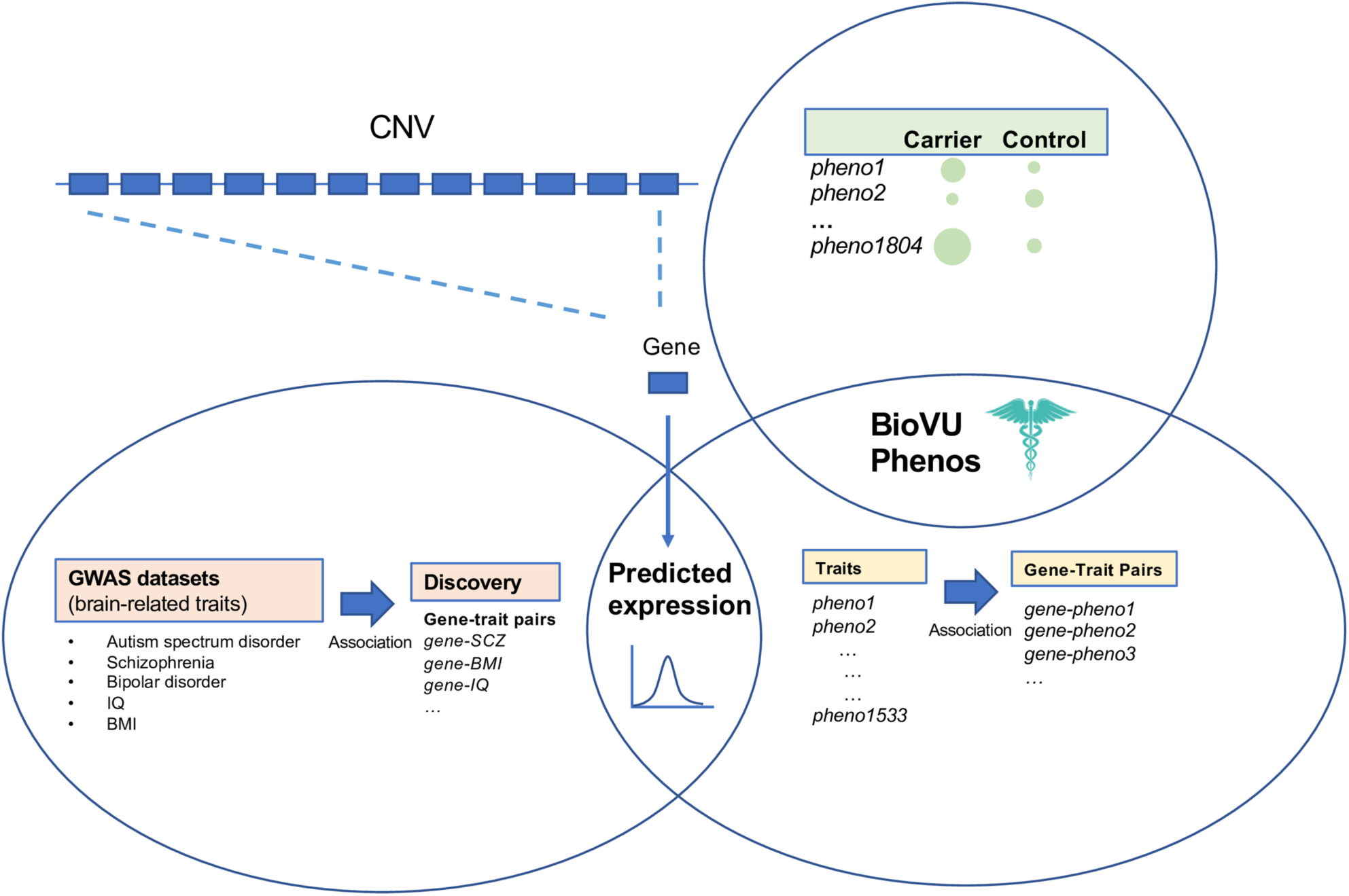
An overview of the three components of this study. We probed the effects of individual genes in the 16p11.2 and 22q11.2 CNVs on phenotype in two ways. First (bottom left), we used large GWAS datasets for brain-related traits associated with both CNVs to determine whether variation in predicted expression in any of the individual genes in each CNV was associated with case-control status for each trait. In the second component of this study (top right), we used a biobank containing clinical and genotypic data to identify the individuals with 16p11.2 and 22q11.2 duplications or deletions and determined the clinical traits that were over-represented in CNV-carriers. Third (bottom right), we used the biobank to perform a phenome-wide association study to determine clinical traits that are driven by the predicted expression of individual CNV genes, as well as whether these traits overlapped with traits over-represented in CNV carriers. Analyses one and three are integrated in their use of imputed expression; analyses two and three are integrated in their use of electronic health data.

## METHODS

### *GWAS Data for* schizophrenia, IQ, BMI, bipolar disorder, and ASD

We obtained the imputed individual-level genotypes for ASD, bipolar disorder, and schizophrenia from the Psychiatric Genomics Consortium in PLINK format (Table S1). These datasets include mainly European populations and are comprised of several independent cohorts: 30 in bipolar disorder (N = 19,202 cases 30,472 controls, downloaded July 2019), 46 in schizophrenia (N = 31,507 cases 40,230 controls, downloaded July 2018), 14 in ASD (N = 7,386 cases, 8,566 controls, downloaded May 2019) ^61, 62, 66^. For two additional traits, we used publicly available summary statistics: BMI from the Genetic Investigation of ANthropometric Traits (GIANT) consortium (2015, both sexes, n=339,224, downloaded June 2019) and IQ from Savage *et al* 2018 (n=269,867, downloaded May 2019) ^63, 64^.

For replication studies and comparison of PheWAS results, we used the publicly available GWAS summary statistics for schizophrenia, IQ, BMI, bipolar disorder, and ASD from the UK Biobank (http://www.nealelab.is/uk-biobank). We could not use the UK Biobank IQ data for replication of our discovery IQ data, as the datasets overlap. The list of UK Biobank phenotypes used is in Table S1. In addition, we used individual-level data from the UK Biobank (n = 408,375) to perform conditional analysis for BMI fine-mapping, but chose not to use it for discovery analysis because of previously-observed high inflation of summary statistics ^67^.

### Expression prediction models

In order to impute gene expression, we obtained PrediXcan models for 48 tissues based on GTEx v7 Europeans from predictdb.org ^58, 59, 68^. These models were generated by training an elastic net model that finds the best set of cis-SNP predictors for the expression of a gene in a tissue in the GTEx cohorts ^58^. Only models with predicted-observed correlation R^2^ > 0.01 and cross-validated prediction performance *P* < 0.05 are kept.

### Genes studied

We studied all coding and noncoding genes at the 16p11.2 and 22q11.2 copy number variant loci for which expression prediction models were available. We included flanking genes in a 200kb window upstream and downstream of the CNV breakpoints. Not all genes in the CNV regions were available to be analyzed through our methods; noncoding genes were especially unlikely to have a high-quality predictive model in any tissue. Thirty-one genes at or near 16p11.2 lacked high-quality prediction models in every tissue. Overall, 46 coding and noncoding genes at or near 16p11.2 were tested. Ninety-nine genes at or near 22q11.2 lacked high-quality prediction models in every tissue. Overall, 83 coding and noncoding genes at or near 22q11.2 were tested (Table S2, Figure S1).

### Comparison of observed expression correlations with predicted expression correlations

Observed expression correlations were calculated at a tissue-specific level on data from GTEx v7 ^69^. Tissue-specific predicted expression was calculated by applying the appropriate GTEx predictive model on the GTEx v6p genotypes (dbgap id: phs000424.v6.p1) for 450 individuals. To minimize spurious correlations, the predicted expression levels were rigorously filtered and normalized. Specifically, the expression levels were filtered for outliers (values above 1.5*interquartile range, in either direction), adjusted for the principal components of both the predicted expression levels and the first 20 PCs of the GTEx genotypes, inverse-quantile normalized, re-adjusted for principal components, and re-filtered for outliers. We observed that normalization of the predicted expression reintroduced correlation between expression and the genotypic PCs, leading us to perform the correction twice.

### Association analysis in individual-level data

All PGC individual-level data analysis was performed on the LISA cluster, part of SURFSara. Each of the three PGC collections went through quality control, filtering, and PCA calculation, as described previously ^60–62^. For each cohort, the convert_plink_to_dosage.py script in PrediXcan was used to convert chromosome 16 and 22 genotypes from PLINK to dosage format, keeping only the SNPs used in at least one predictive model. Using these dosages, the --predict function in PrediXcan was used to generate predicted expressions of CNV genes for each individual. Genes with predicted expression of 0 for all individuals in a single tissue were filtered out. The average number of genes filtered out across tissues and cohorts was 0.89; the maximum was 11 in Thyroid in the Japanese cohort. Cross-tissue association studies between predicted expression and case-control status were performed using MultiXcan. In brief, MultiXcan takes the matrix of predicted expressions across tissues for a single gene, calculates the principal components of this matrix to adjust for collinearity, fits a model between these PCs and case-control status, and reports the results of the overall fit ^59^. As in the PGC association studies, our analysis was adjusted by the principal components that were significantly associated with each trait– 7 for bipolar disorder, 10 for schizophrenia, and 8 for autism case-control studies (the autism trios were not adjusted for covariates). UK Biobank MultiXcan analysis was limited to white ethnicity, and included age, age-squared, 40 principal components as covariates.

Meta-analysis with METAL on the p-values from MultiXcan, weighted by the sample size of each cohort, was used to calculate a cross-cohort association statistic ^70^. The joint fit in MultiXcan generates an F-statistic that is always greater than zero, while some of our traits of interest have a specific expected direction (only in deletion or only a duplication). Thus, a direction was assigned to each MultiXcan result. This was done by running a tissue-specific PrediXcan association analysis between predicted expressions and case-control status (using -- logistic), which calculates a signed association Z-score for every gene. The sign of the mean Z-score for that gene across all tissues was the direction of association used for meta-analysis.

### Association analysis in summary-level data

Both the single-tissue PrediXcan and the multi-tissue MultiXcan methods have been extended to estimate the association results between genetically regulated expression and a trait if only the summary statistics for the trait are available. For each trait’s summary statistics, the summary version of PrediXcan (S-PrediXcan) and the associated MetaMany.py script was used to calculate the per-tissue association results for each gene in 48 GTEx tissues. Association results were aggregated across tissues using the summary version of MultiXcan (S-MultiXcan). The mean single-tissue Z-score (as reported in the zmean column in the S-MultiXcan output) was used as the direction of association. The UK Biobank replication studies were performed in the same way.

### Conditional analysis to fine-map associations

In order to adapt the multi-tissue association analysis to perform conditional testing, “conditioned predicted expressions” were generated for a set of genes associated with the same trait. As an example, take the set of three genes [*INO80E, YPEL3, TMEM219*] associated with schizophrenia. In order to condition on *INO80E*, for example, the predicted expression of *INO80E* was regressed out of the predicted expressions of *YPEL3* and *TMEM219*. Conditioning was only done in tissues where the predicted expressions of the genes were correlated (Spearman correlation *P* < 0.05). Another set of conditioned predicted expressions was generated by adjusting the predicted expression of *INO80E* by the predicted expressions of [*TMEM219, YPEL3*]. Separately, these per-tissue conditioned predicted expressions were used as inputs for a MultiXcan analysis and METAL meta-analysis on schizophrenia as described earlier. All three individually associated genes were tested in this manner. The same analysis was later used to test for independence of association between BMI in the UK Biobank as well as *psychosis* and *morbid obesity* traits in the PheWAS. The *P_cond_* reported in the text is the p-value of a gene-trait pair when adjusting for all other genes considered for conditioning for this trait, unless otherwise stated.

### Phenome-wide association studies

Vanderbilt University Medical Center (VUMC) houses de-identified phenotypic data in the form of the electronic health records (EHR) within the synthetic derivative (SD) system ^71^. The SD contains EHR data including ICD9/10 billing codes, physician notes, lab results, and similar documentation for 3.1 million individuals. BioVU is a biobank at VUMC that is composed of a subset of individuals from the SD that have de-identified DNA samples linked to their EHR phenotype information. The clinical information is updated every 1-3 months for the de-identified EHRs. Detailed description of program operations, ethical considerations, and continuing oversight and patient engagement have been published ^71^. At time of analysis, the biobank contained 48,725 individuals who had been genotyped. DNA samples were genotyped with genome-wide arrays including the Multi-Ethnic Global (MEGA) array, and the genotype data were imputed into the HRC reference panel ^72^ using the Michigan imputation server ^73^. Imputed data and the 1000 Genome Project data were combined to carry out principal component analysis (PCA) and European ancestry samples were extracted for analysis based on the PCA plot. GTEx v7 models from PredictDB were applied to the samples to calculate genetically regulated expression (GReX).

Phenome-wide association studies (PheWAS) were carried out using ‘phecodes’, phenotypes derived from the International Code for Diseases version 9 (ICD-9) billing codes of EHRs. The PheWAS package for R, version 0.11.2-3 (2017) was used to define case, control and exclusion criteria ^74, 75^. We required two codes on different visit days to define a case for all conditions, and only phecodes with at least 20 cases were used for analysis (1531 traits). The single-tissue predicted expressions were combined across tissues using MultiXcan, as was done to analyze individual-level GWAS data from the Psychiatric Genomics Consortium ^59^. Covariates for this analysis were age, sex, genotyping array type/batch and three principal components of ancestry.

The top 1% (top 15 traits) of every gene’s association results was kept for analysis. A binomial test was used to compare whether the number of traits in any clinical category (circulatory system, genitourinary, endocrine/metabolic, digestive, neoplasms, musculoskeletal, injuries & poisonings, mental disorders, sense organs, neurological, respiratory, infectious diseases, hematopoietic, symptoms, dermatologic, congenital anomalies, pregnancy complications) were over-represented in the top 1% of results compared to the proportion of each category among all 1531 traits tested. The expected number of each clinical category as determined by [15 traits * n_genes_]*p_i_ where p_i_ is the probability of a randomly drawn (without replacement) code belongs to category i. P_i_ can be estimated by the number of codes belonging to category i divided by all codes tested (n=1531). The significance threshold was 0.05/[17 categories] = 0.0029.

### Determining traits over-represented in carriers

3.1 million electronic medical records from the SD at VUMC were queried for keywords corresponding to copy number variations at 16p11.2 and 22q11.2 (Table S3). Individual charts identified as containing the keywords were manually reviewed and patients were labeled as cases if their medical records provided evidence of CNV carrier status. Patients identified in the queries with insufficient evidence of CNV carrier status were excluded from the analysis. Cases with positive 16p11.2 and 22q11.2 CNV carrier status were identified as: “16p11.2 duplication” (n=48, median age 11), “16p11.2 deletion” (n=48, median age 12), “22q11.2 duplication” (n=43, median age 11). Additional individuals in the 22q11.2 deletion case group were identified by querying the medical records for alternate terms including: “velocardiofacial”, “DiGeorge”, “conotruncal anomaly face”, “Cayler”, “Opitz G/BBB”, “Shprintzen”, and “CATCH22” (n=388, average age 17). Individuals were excluded from case groups if they were included in the genotyped sample used for the gene-by-gene analysis, or if their records included a mention of additional CNVs. Individuals within the 16p11.2 case groups were also excluded if the size of the reported CNV was 200-250 kb. Individuals within the 22q11.2 case group were excluded if the size of the CNV was smaller than 500 kb or if there was a mention of “distal” when referring to the deletion or duplication. PheWAS was carried out, with each of the four carrier categories as cases and over 700,000 medical home individuals as controls, using age, sex, and self-reported race as covariates. The medical home individuals are patients seen at a Vanderbilt affiliated clinic on five different occasions over the course of three years. Because the sample size for this analysis was larger (700,000 individuals vs. 48,000), and we used traits only that were present in 20 or more individuals, there were more traits available for analysis here, n=1795. After calculating PheWAS, we excluded over-represented traits that were present in <5% of carriers from further analyses.

### Test for enrichment of gene-specific PheWAS for traits over-represented in carriers

For each of 16p11.2 duplications, 16p11.2 deletions, 22q11.2 duplications, 22q11.2 deletions, the entire carrier vs. non-carrier PheWAS results were ranked. All of the traits in the top 1% of per-gene 16p11.2 and 22q11.2 PheWAS results were converted to a value corresponding to the rank of the trait in the carrier vs. non-carrier PheWAS. To determine whether the CNV genes’ PheWAS top traits were distributed nonrandomly within the ranks of all traits, the distribution of the ranks of the each CNV’s per-gene PheWAS top traits was compared to the ranks of all carrier vs. noncarrier PheWAS traits for the same CNV (a uniform distribution) using a Kolmogorov-Smirnov test.

### Significance threshold for association studies

The significance threshold used for each discovery MultiXcan or S-MultiXcan association study and conditional analysis was 0.05/(number of traits*number of CNV genes tested). In practice, this usually meant 5 traits and 132 CNV genes, for a threshold of *P* < 7.6×10^-5^. For replication studies, the significance threshold was set at 0.05 in order to test a single gene. The exception was in the BMI UK Biobank dataset. We first tried a phenotype-swapping approach to generate an expected distribution for the 16p11.2 genes. The distributions were null and did not yield meaningful comparisons. Instead, 100 random subsets of adjacent genes of approximately the same length and gene count as the CNV were tested for association with BMI. The 95th percentile of the MultiXcan p-values for these genes was used as a permutation-based significance threshold.

In the gene-based PheWAS study, there were 1,534 phecodes (each with at least 20 cases) tested overall, corresponding to a Bonferroni-corrected phenome-wide significance threshold of 3.3×10^-5^. For genes having no phenome-wide significant results, their top 15 associations, corresponding to the top 1% of the 1,534 phecodes, were used. In the carrier vs. non-carrier PheWAS, there were 1,795 phecodes tested overall, corresponding to a Bonferroni-corrected phenome-wide significance threshold of 2.79×10^-5^. Additional traits meeting a false discovery rate threshold of 0.05 were considered in identifying traits both over-represented in carriers and represented in individual gene PheWAS.

### Graphical summary of selected PheWAS results

The *chordDiagram* method in the *circlize* package was used to generate the circle summary plots ^76^. The gene-trait pairs we selected for Tables 1 and 2 were used as inputs, with the -log10 p-value of association used as the weighting to determine the edge width. For the 22q11.2 circle plot, only associations with *P* < 5×10^-3^ were used in order to create a legible plot. Descriptions were cut off at 55 characters; to read the entire descriptions see Tables 1 and 2.

**Table 1.**
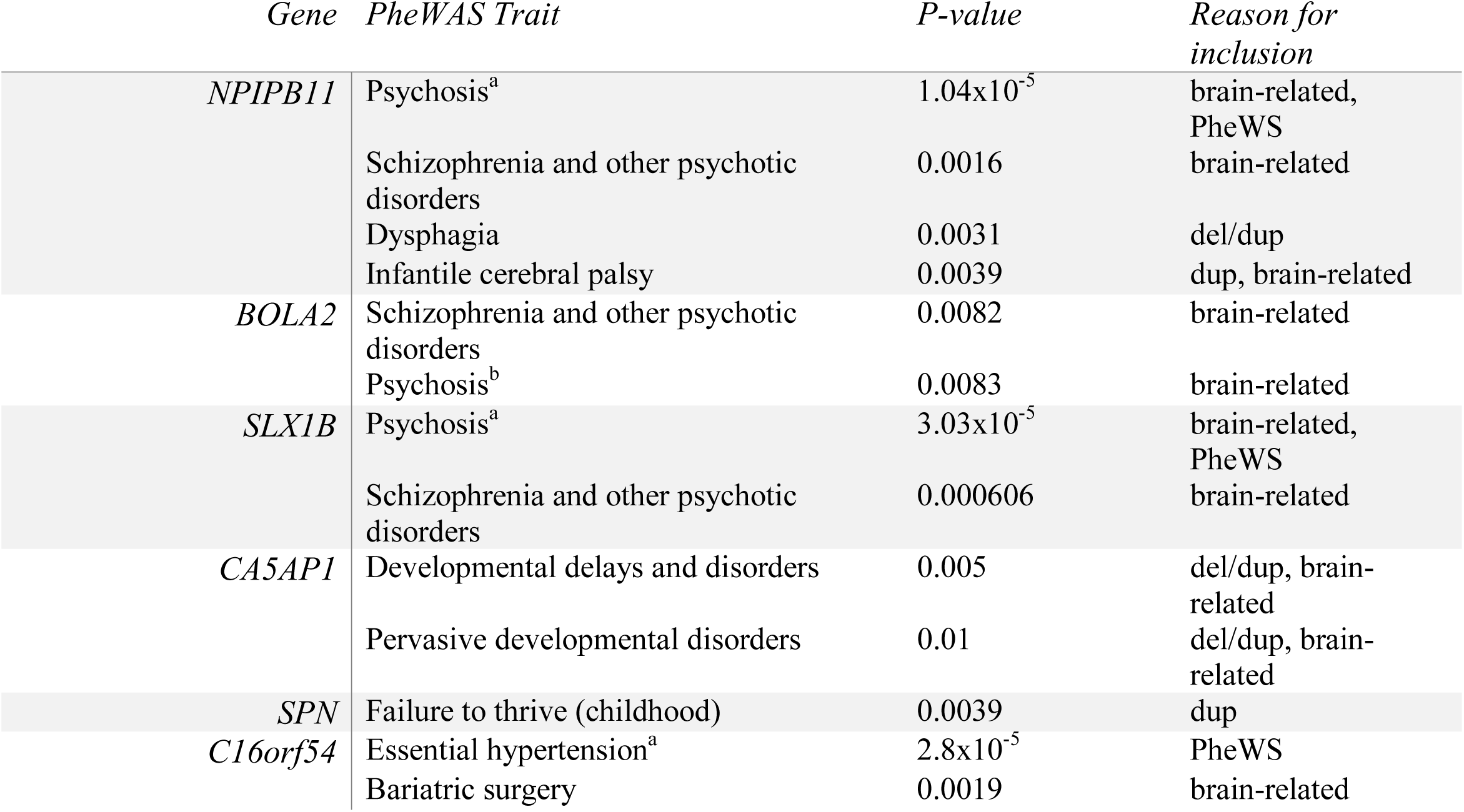
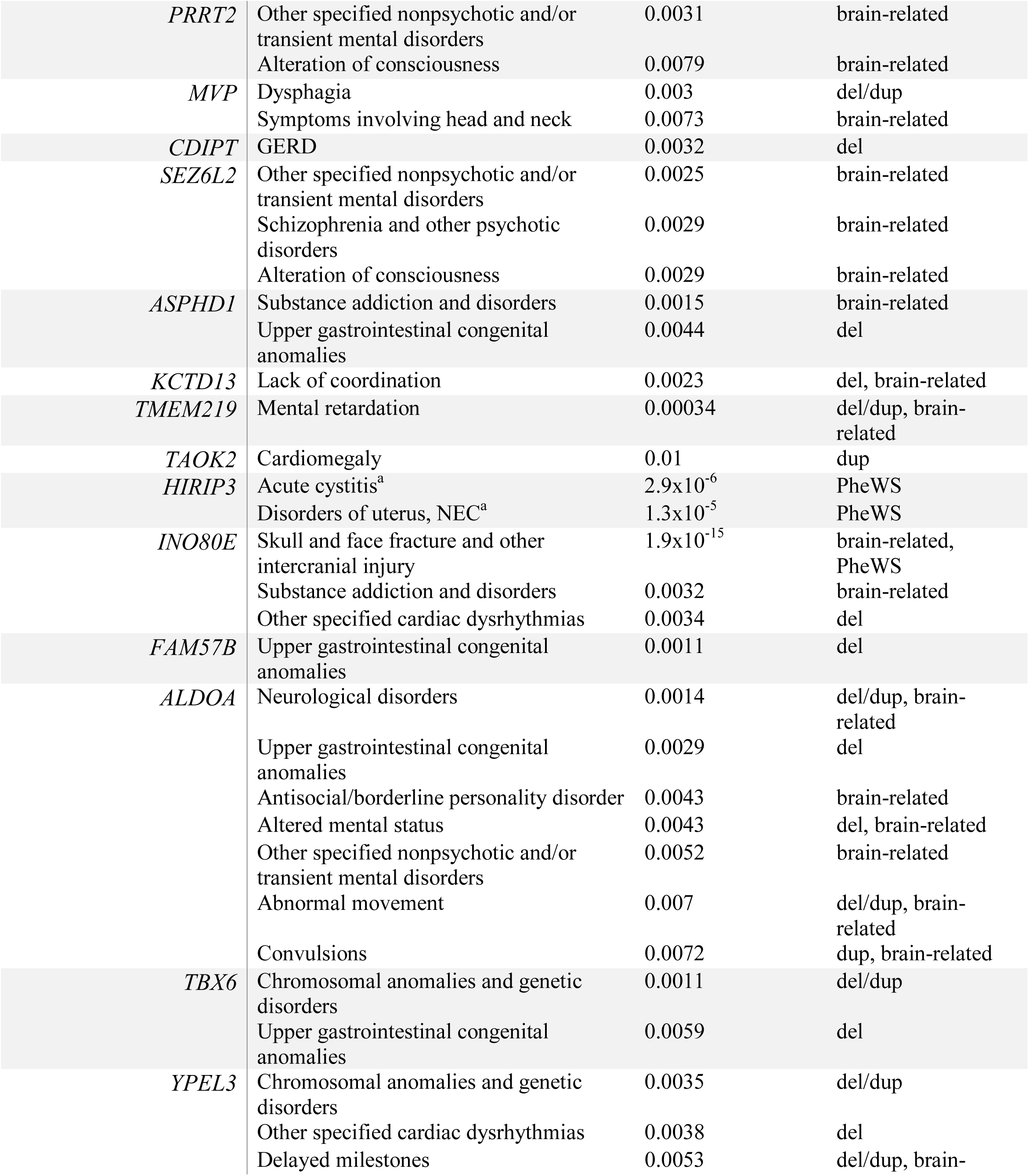
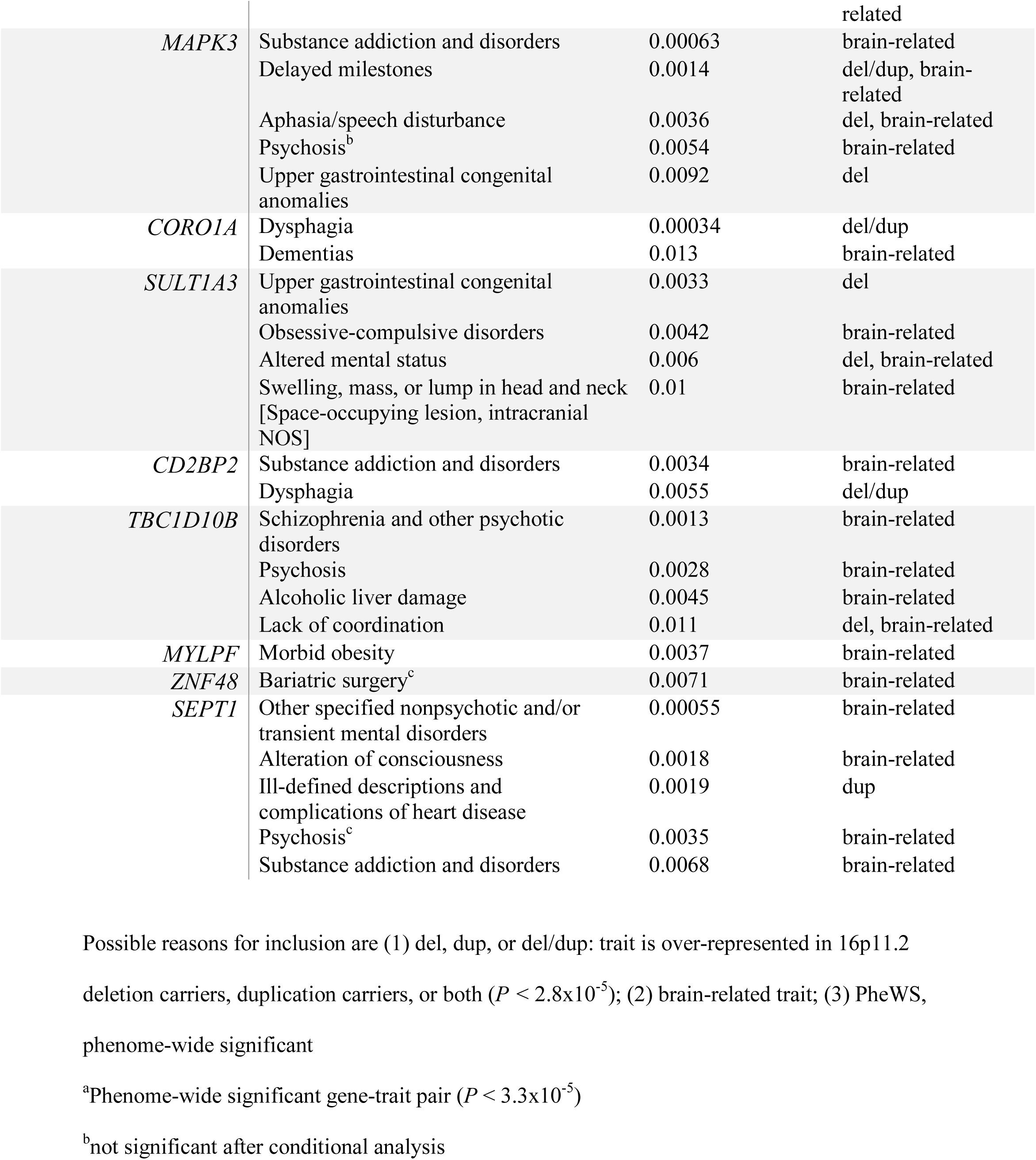
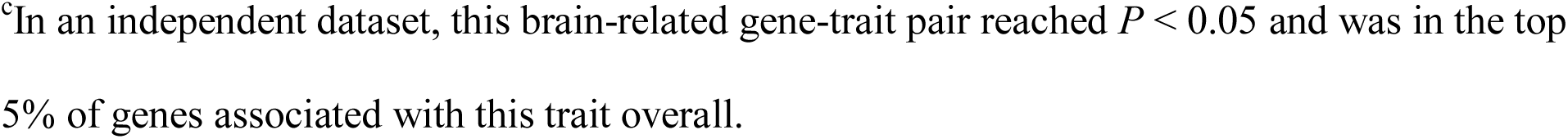
Selected 16p11.2 gene associations with PheWAS traits

**Table 2.**
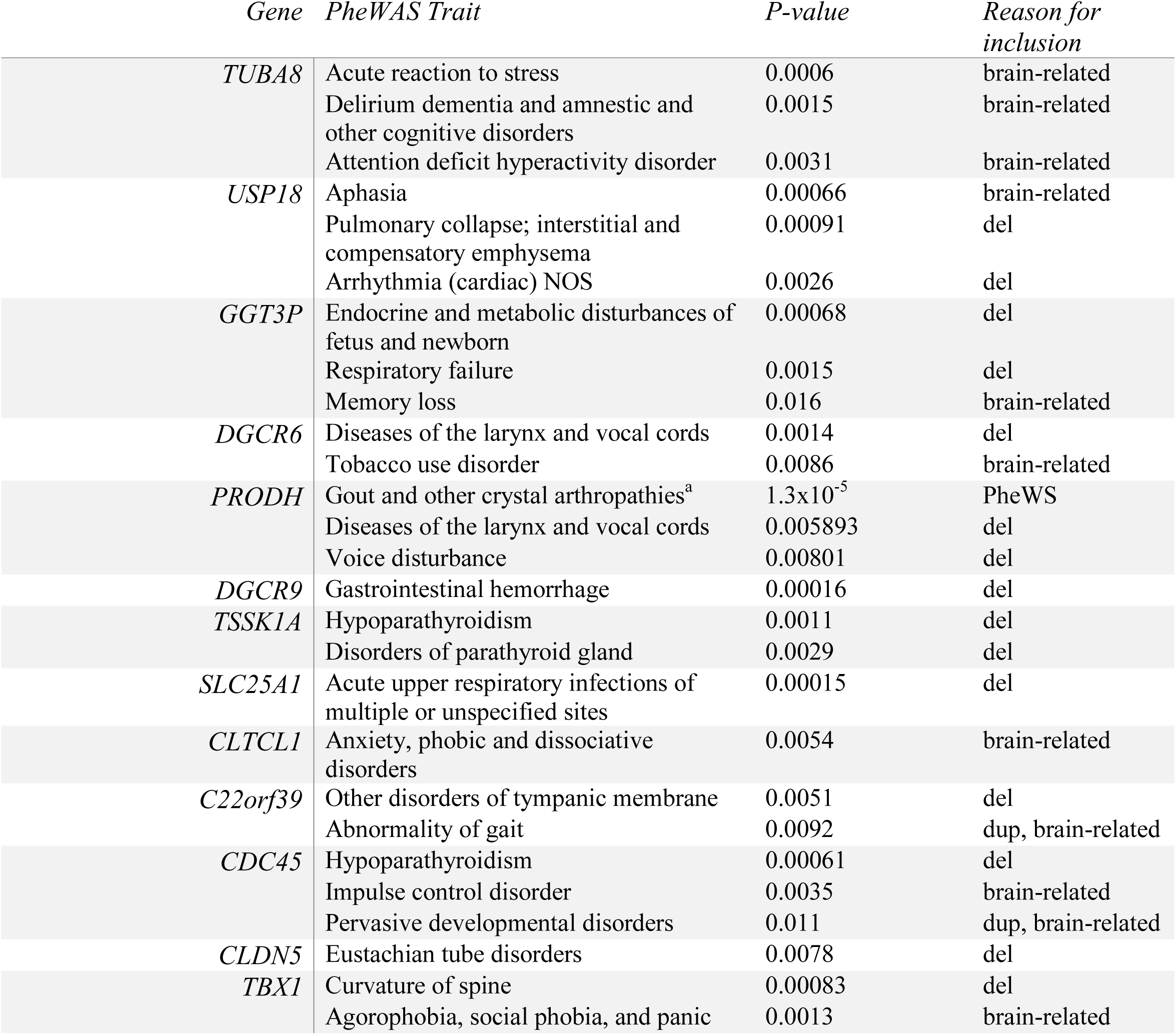
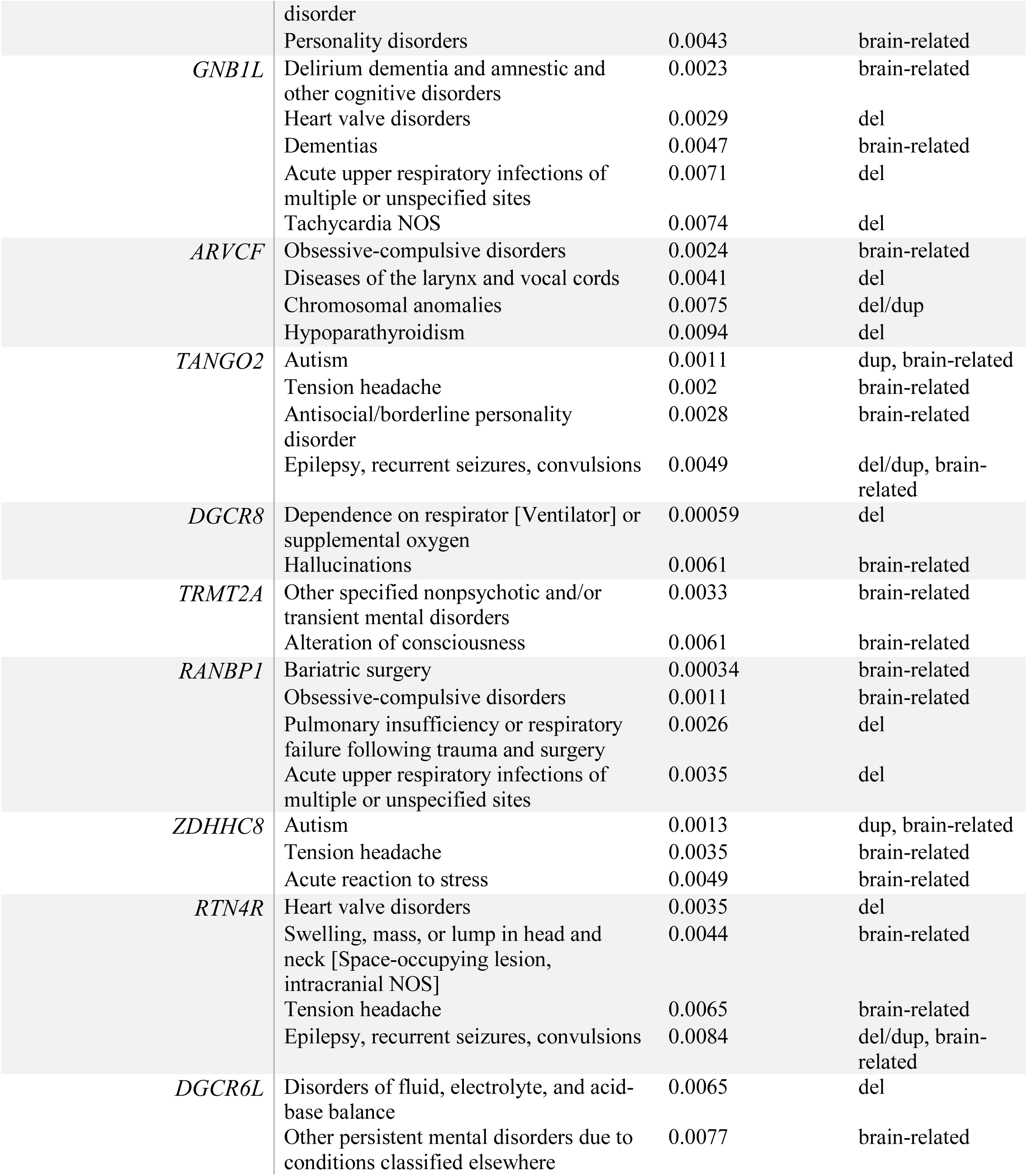
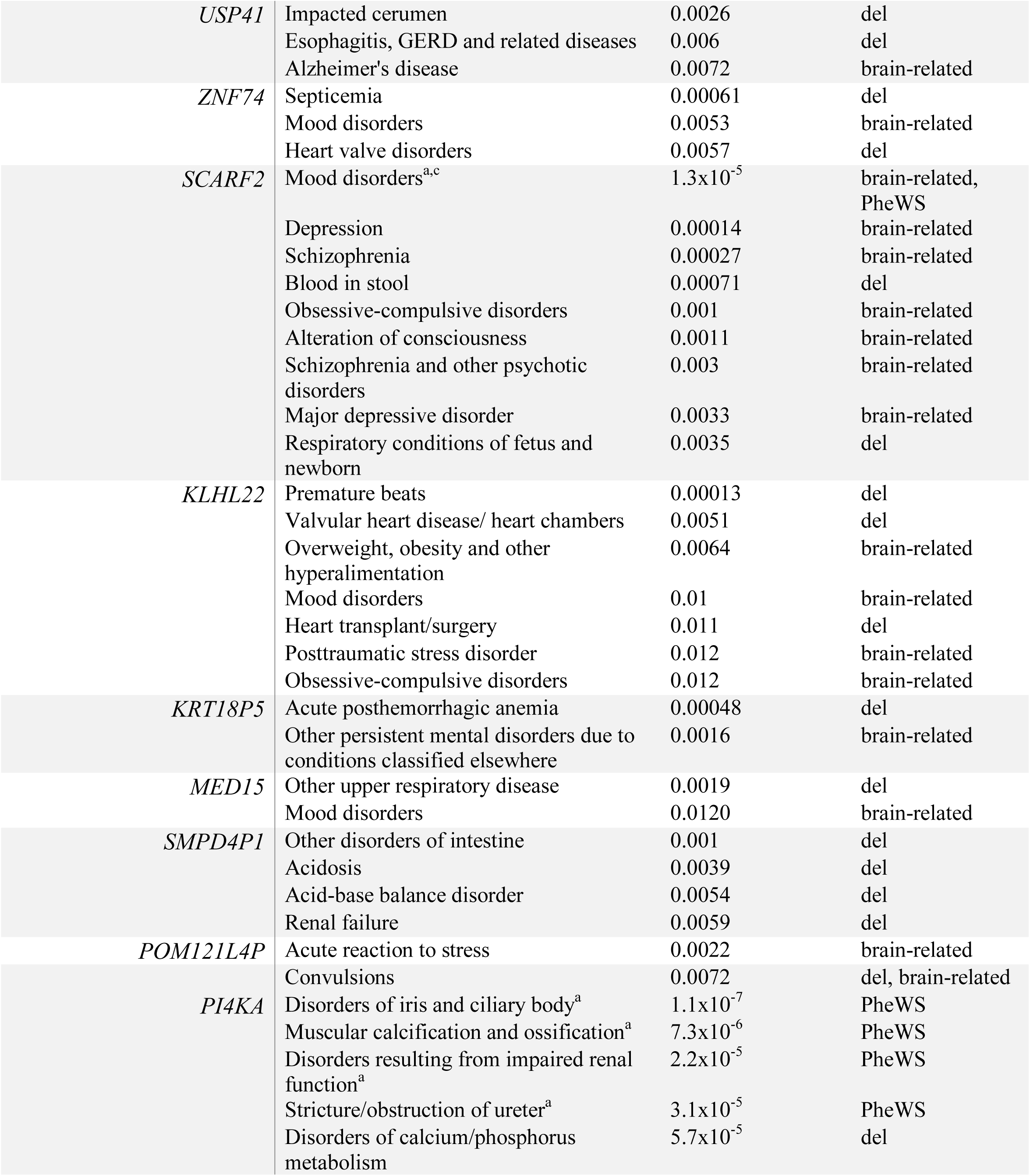
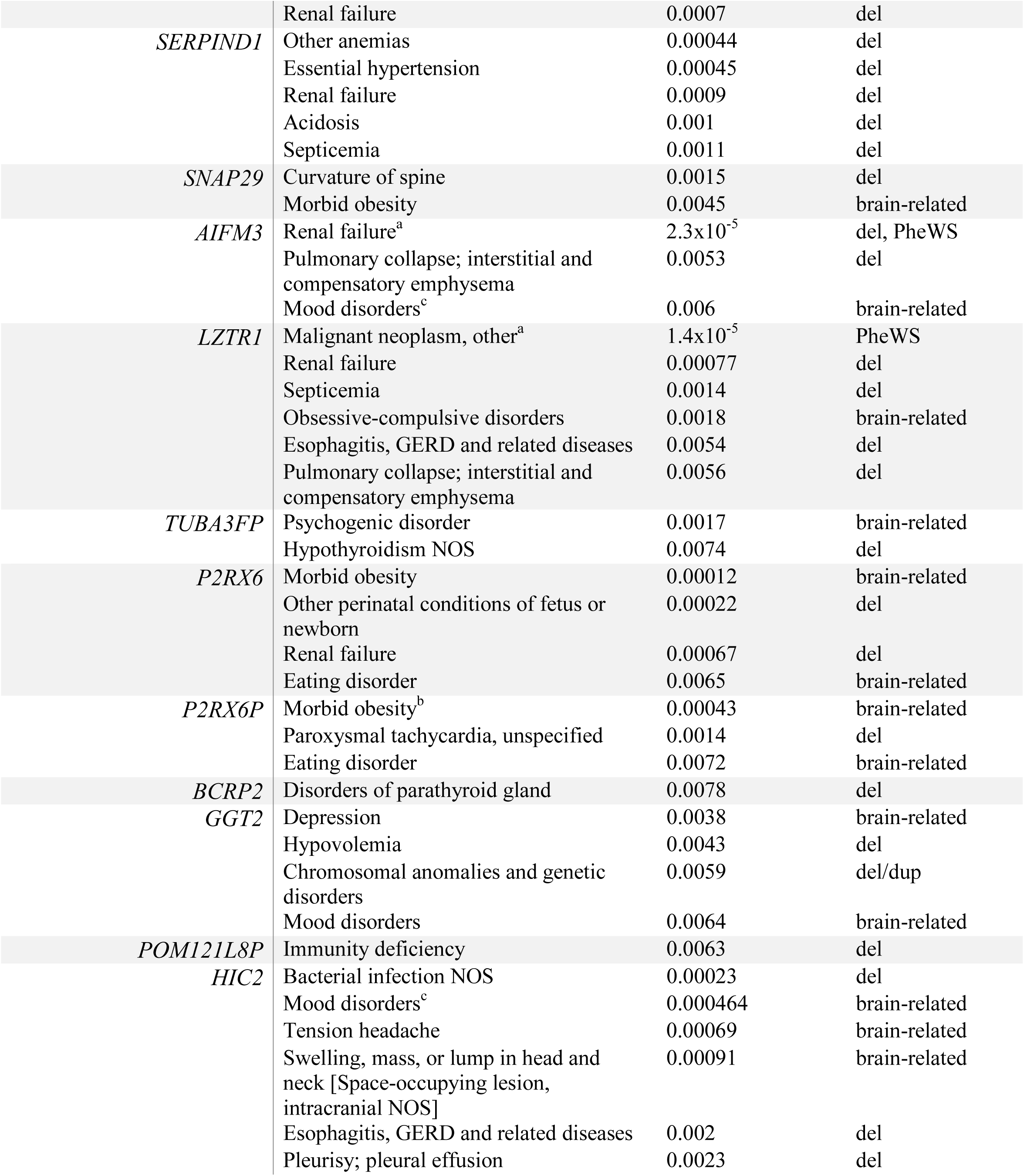
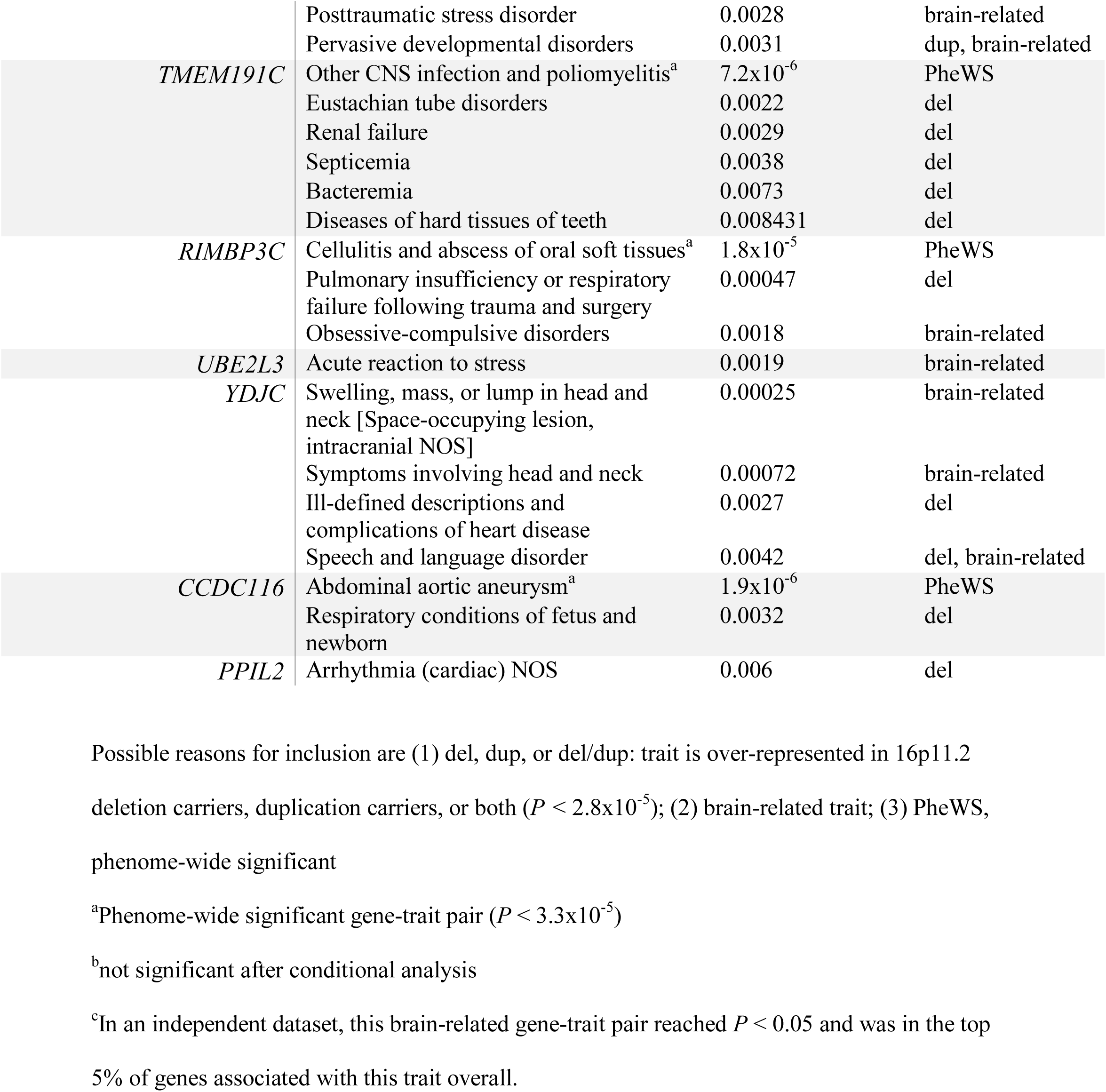
Selected 22q11.2 gene associations with PheWAS traits

## RESULTS

### Individual genes at 16p11.2 are associated with schizophrenia, IQ, and BMI

In order to find genes at copy number variant loci driving brain-related disorders, we performed an association analysis between imputed gene expression levels and five traits: schizophrenia, IQ, BMI, bipolar disorder, and ASD. It has been observed that the 16p11.2 and 22q11.2 copy number variants affect expression of nearby genes, so we included genes flanking the CNV locus by 200kb in both directions ^45, 46, 77^. Overall, we tested 83 coding and noncoding genes at or near 22q11.2 and 46 genes at or near 16p11.2 for which a predictive model was available (Table S2, Figure S1). As cis-eQTLs are often shared among tissues, we pooled together information from all tissues in GTEx to boost our power to detect brain-related traits ^59^.

Two genes at 16p11.2 show predicted expression positively associated (*P* < 7.6×10^-5^) with schizophrenia (Figure 2, S3 Table): *TMEM219* (*P* = 1.5×10^-5^) and *INO80E* (*P* = 5.3×10^-10^). This positive direction of effect is consistent with the association between 16p11.2 duplications and schizophrenia ^2^. An additional gene, *YPEL3*, was significantly associated with schizophrenia in the negative direction (*P* = 4.9×10^-6^). For IQ, there was one strong positive association at the 16p11.2 locus (Figure 2, S3 Table): *SPN* (*P* = 2.9×10^-22^). Intellectual disability is observed in both deletions and duplications of 16p11.2, so there was no expected direction of effect ^3,14^. Four genes showed negative association with BMI (Figure 2, S3 Table): *SPN* (*P* = 6.2×10^-18^), *TMEM219* (*P* = 2.2×10^-5^), *TAOK2* (*P* = 8.5×10^-11^), and *INO80E* (*P* = 1.0×10^-7^). We focused on genes with negative associations with BMI because, in humans, obesity is associated with deletions at 16p11.2 ^1,17^. Two additional genes, *KCTD13* (*P* = 9.5×10^-6^) and *MVP* (*P* = 2.1×10^-5^), were significantly associated with BMI in the positive direction. No gene at 16p11.2 was significantly associated with bipolar disorder or ASD (Table S4, Figure S2). No individual genes at or near 22q11.2 had predicted expression significantly associated with any of the five traits (Table S4, Figure S3).

**Figure 2.**
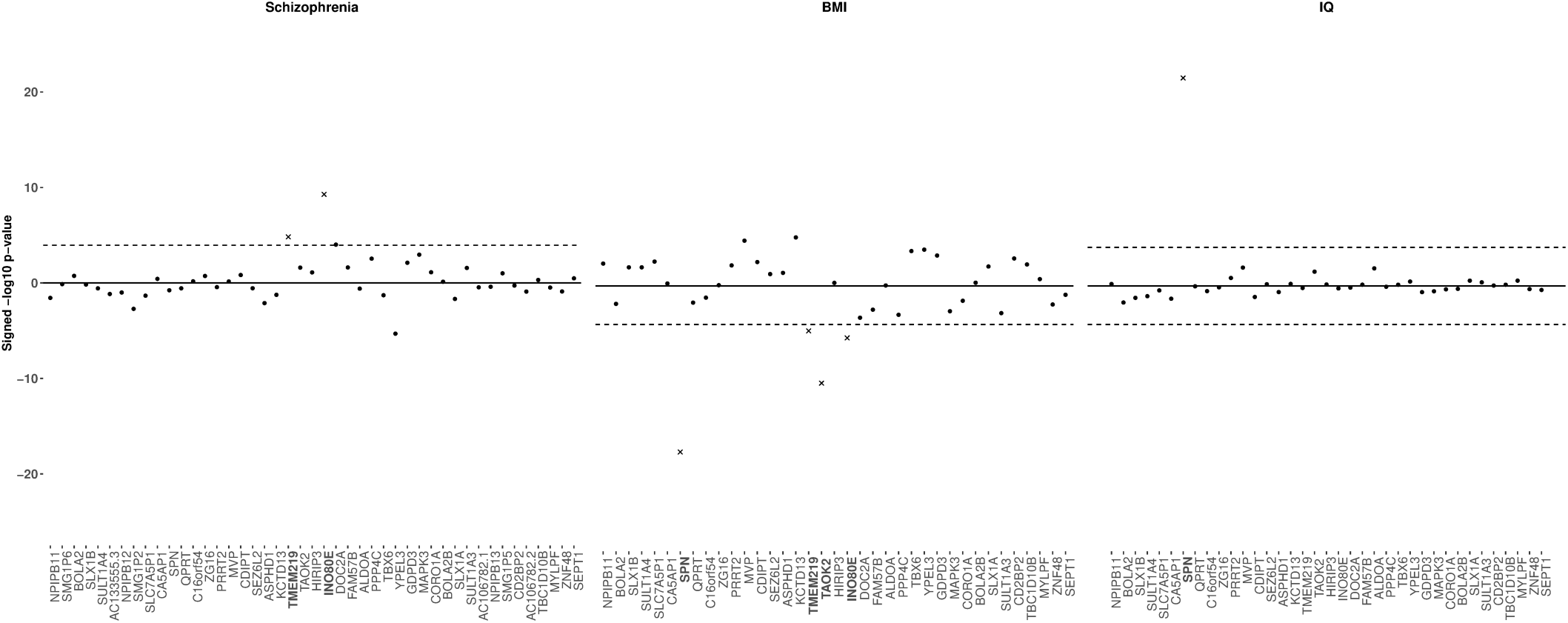
Association between 16p11.2 genes and three brain-related traits. Association between predicted expression of 16p11.2 genes and schizophrenia (left), BMI (middle), IQ (right) using MultiXcan (schizophrenia) and S-MultiXcan (BMI, IQ). Genes are listed on the horizontal access in order of chromosomal position. The -log10 p-values on the vertical axis are given a positive or negative direction based on the average direction of the single-tissue results. The significance threshold, *P* < 7.6×10^-5^, is a Bonferroni correction on the total number of 16p11.2 and 22q11.2 genes (132) tested across 5 traits (0.05/(5*132)). Genes exceeding the significance threshold in the expected direction (positive for schizophrenia, negative for BMI, either for IQ) are denoted as x’s.

### Follow-up conditional analyses narrow down genes driving schizophrenia and BMI

To replicate our analysis, we used a large UK Biobank cohort for which GWAS summary statistics were available for multiple brain-related traits (Table S1) ^78^. The predicted expression of *INO80E* and *TMEM219* from the discovery analyses were associated (*P* < 0.05) with having an ICD10 diagnosis of schizophrenia (ICD10: F20, 198 cases: *INO80E P* = 0.04, *TMEM219 P =* 0.03, Table S6). Although this is only nominally significant, it is notable that these genes are in the 3^rd^ percentile of schizophrenia associations genome-wide within UK Biobank.

The UK Biobank GWAS of BMI is highly inflated, including in the 16p11.2 region. Nearly every 16p11.2 gene showed association at the previously used threshold (*P* < 7.6×10^-5^). Using a permutation-based approach within individual-level data, we adjusted the significance threshold to 8.8×10^-11^. All genes from the discovery analysis replicated (Table S6): *SPN* (*P* = 6.1×10^-23^), *KCTD13* (*P =* 1.2×10^-30^), *TMEM219* (*P =* 7.1×10^-37^), *MVP* (*P =* 5.1×10^-11^) *INO80E (P* = 1.9×10^-27^). We were not able to replicate the IQ result in the UK Biobank, because the UK Biobank sample overlapped with our discovery GWAS.

We performed an additional fine-mapping study on the three genes associated with schizophrenia. Linkage disequilibrium between the eQTL SNPs in predictive models may lead to correlation among predicted expressions for nearby genes, so it is possible not all three detected association signals are independent. The predicted expressions of *INO80E, YPEL3*, and *TMEM219* were moderately correlated (the correlation of *INO80E* to the other genes is in the range of -0.4 to 0.37 across GTEx tissues, for example), consistent with the relationships between the observed expressions of these genes (measured expression of *INO80E* is correlated to measured expression of the other genes in the range -0.36 to 0.31). In order to pick out the gene(s) driving the association signal, we used a conditional analysis approach (Table S5). We observed that after adjusting the predicted expression of the other CNV genes for the predicted expression of *INO80E*, no gene was significantly associated with schizophrenia. However, when we adjusted the predicted expression of *INO80E* by the predicted expressions of the other two highly associated genes, *INO80E* remained significantly associated with schizophrenia (*P* = 2.3×10^-6^). The same pattern was not observed for *TMEM219* or *YPEL3*, suggesting *INO80E* explains the entire 16p11.2 signal for schizophrenia.

While we did not have individual level data for the GIANT consortium, we obtained individual-level BMI data from the UK Biobank. We performed an analogous conditional analysis on the six genes associated with BMI, *SPN, INO80E, TMEM219, TAOK2* in the negative direction, as well as *KCTD13* and *MVP* in the positive direction. Due to the inflation in the UK Biobank data, all of these genes had very low p-values even after conditioning; however, we see that some genes’ association results stayed in the same range, while others increased in p-value by five orders of magnitude or more after adjusting by the other five genes. Based on these observations, it is likely that *SPN* (*P_UKBB_ =* 6.1×10^-23^, *P_cond_ =* 7.5×10^-21^), *INO80E* (*P_UKBB_* = 1.9×10^-27^, *P_cond_ =* 2.8×10^-32^), and *KCTD13* (*P_UKBB_ =* 1.2×10^-30^, *P_cond_ =* 4×10^-27^) were independently associated with BMI, while *TMEM219* (*P_UKBB_ =* 7×10^-37^, *P_cond_ =* 2.3×10^-18^)*, TAOK2* (*P_UKBB_ =* 4.2×10^-29^, *P_cond_ =* 2.3×10^-19^), and *MVP* (*P_UKBB_ =* 5.1×10^-11^, *P_cond_* = 5×10^-6^) were significant in the discovery analysis primarily due to correlation with one of the independent genes.

### Phenome-wide association studies identify previously known and novel traits associated with 16p11.2 and 22q11.2 carrier status

While GWAS datasets provide insight into the impact of genes on ascertained brain-related traits, the 16p11.2 and 22q11.2 CNVs may contribute to a wide spectrum of traits, including milder manifestations of brain-related traits. Thus, biobanks containing both genetic and clinical data can tell us about broader clinical impacts on medical traits. We queried the de-identified electronic health records for 3.1 million patients at VUMC to explore the impacts of the 16p11.2 and 22q11.2 CNVs, as well as their individual genes, on the medical phenome in a representative population ^71^. CNV diagnoses are documented in the medical records, which led us to ask: what are the specific clinical phenotypes that are common in individuals identified as 16p11.2 and 22q11.2 CNV carriers? Carriers were identified by diagnosis of 16p11.2 or 22q11.2 deletion/duplication (or syndromic names for 22q11.2, see methods) in their medical record, and over 700,000 individuals were used as controls. We performed a phenome-wide association study (PheWAS) between 16p11.2 and 22q11.2 deletion/duplication carriers and controls against 1,795 medical phenotype codes (Figs 3 and 4) ^74, 79^. Traits that were significantly over-represented in carriers (*P <* 2.8×10^-5^) fell into three major categories: (1) known primary CNV clinical features, including possible reasons for the referral of the patient for genetic testing (i.e. neurodevelopmental concerns, epilepsy, congenital heart defects), (2) secondary CNV features known to be present in carriers but unlikely to be a primary reason for referral for genetic testing, (3) novel diagnoses not previously reported (Figure 3, Figure 4, Table S7). We chose to focus on traits present in at least 5% of carriers to avoid over-interpreting rare traits.

**Figure 3.**
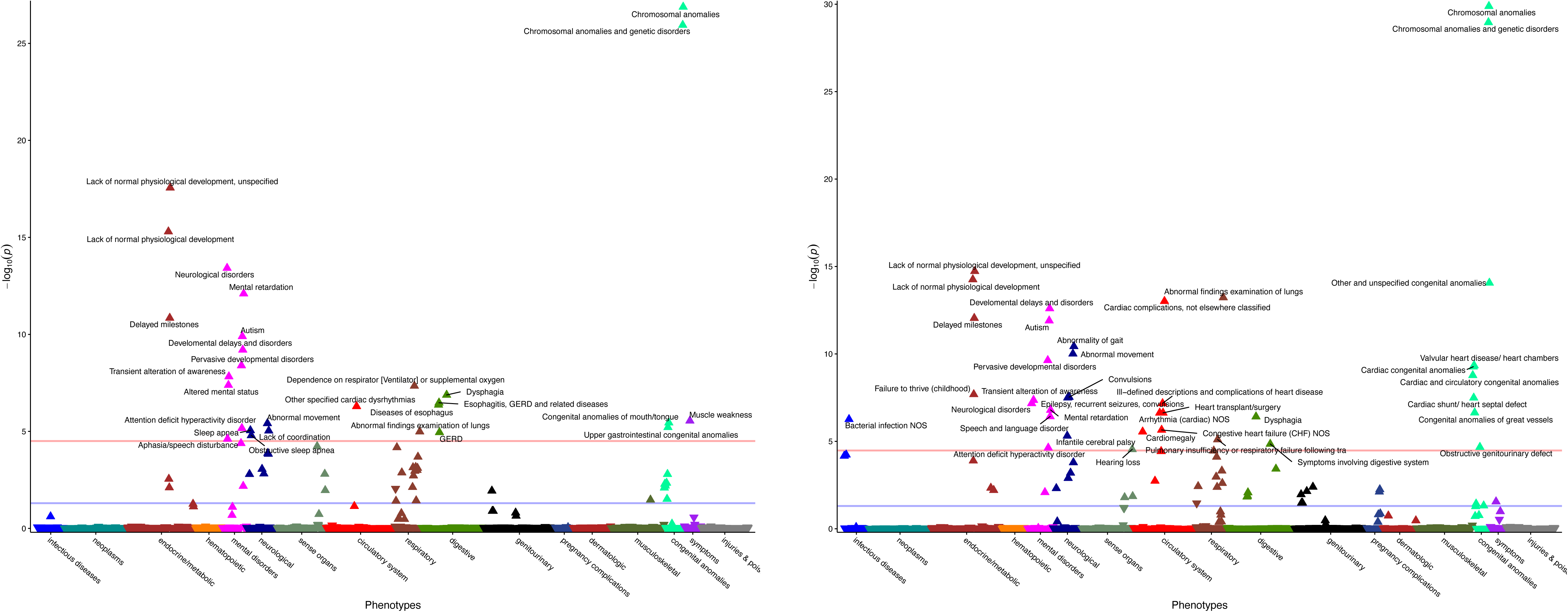
Clinical traits over-represented in 16p11.2 deletion and duplication carriers. CNV carriers were identified in the EHR by keyword search and chart review (left, 16p11.2 deletions [n=48], right, 16p11.2 duplications [n=48], see Methods). Controls included all individuals without the CNV within the medical home population at Vanderbilt (n∼707,000). The x-axis represents the PheWAS codes that are mapped from ICD-9/ICD-10 codes, grouped and color-coded by organ system. The y-axis represents the level of significance (-log_10_*p*). The horizontal red line indicates a Bonferroni correction for the number of phenotypes tested in this PheWAS (*p* = 0.05/1,795 = 2.8×10^-5^); the horizontal blue line indicates *p* = 0.05. Each triangle represents a phenotype. Triangles represent direction of effect; upward pointing arrows indicate phenotypes more common in cases. Covariates included age, sex, and self-reported race extracted from the EHR. Phenotypes reaching Bonferroni-corrected significance level are labeled in plot.

**Figure 4.**
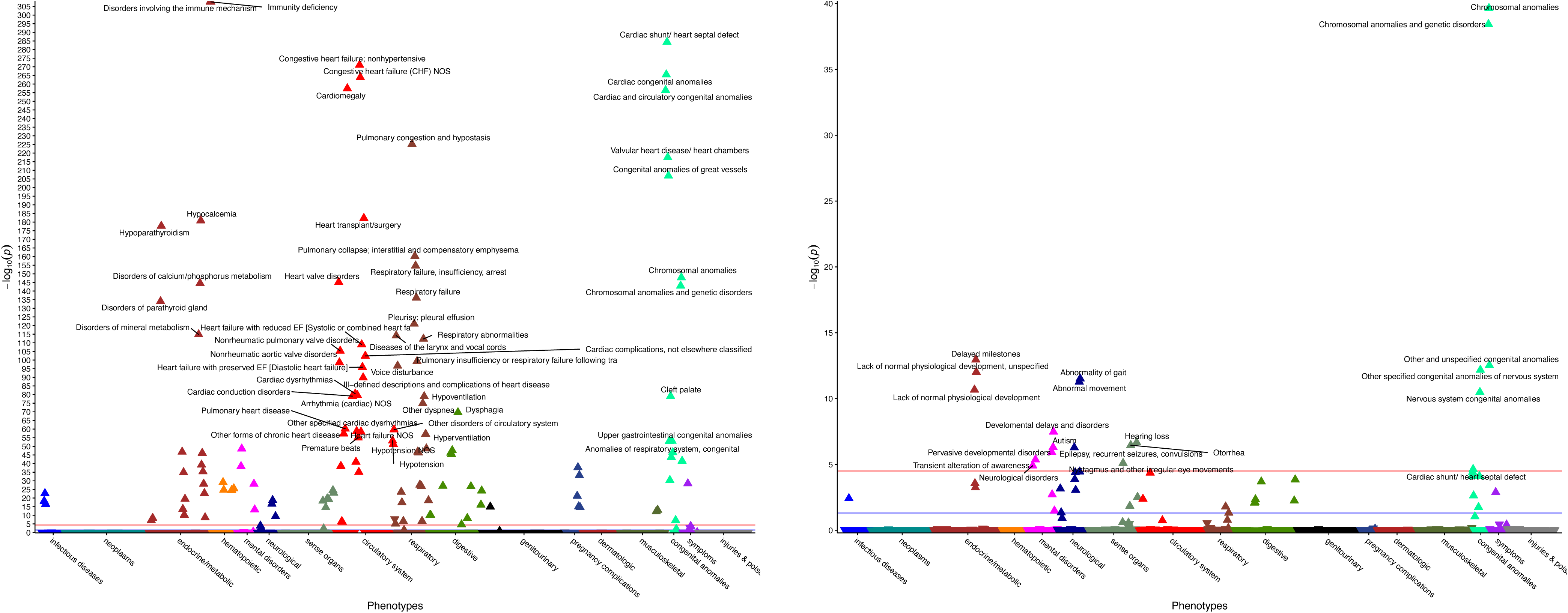
Clinical traits over-represented in 22q11.2 deletion and duplication carriers. CNV carriers were identified in the EHR by keyword search and chart review (left, 22q11.2 deletions [n=388], right, 22q11.2 duplications [n=43], see Methods). Controls included all individuals without the CNV within the medical home population at Vanderbilt (n∼707,000). The x-axis represents the PheWAS codes that are mapped from ICD-9/ICD-10 codes, grouped and color-coded by organ system. The y-axis represents the level of significance (-log_10_*p*). The horizontal red line indicates a Bonferroni correction for the number of phenotypes tested in this PheWAS (*p* = 0.05/1,795 = 2.8×10^-5^); the horizontal blue line indicates *p* = 0.05. Each triangle represents a phenotype. Triangles represent direction of effect; upward pointing arrows indicate phenotypes more common in cases. Covariates included age, sex, and self-reported race extracted from the EHR. Top phenotypes (*P* < 1.0×10^-50^) are labeled in the 22q11.2 deletion plot (left). Phenotypes reaching Bonferroni-corrected significance level are labeled in the 22q11.2 duplication plot (right).

16p11.2 deletion carrier status was associated with developmental diagnoses (Figure 3): *lack of normal physiological development* (*P* = 2.8×10^-18^), *developmental delays and disorders* (*P*= 6.3×10^-10^), *delayed milestones* (*P* = 1.4×10^-11^) ^3^. In addition, 16p11.2 deletion carrier status was associated with *autism* (*P* = 1.3×10^-10^) and *mental retardation* (*P* = 7.9×10^-13^) ^5^. The digestive diagnosis of *GERD* (*P =* 1.1×10^-5^) has been previously observed in carriers but was unlikely to be a primary reason for genetic testing ^80^. *GERD* was accompanied by other digestive diagnoses such as *dysphagia* (*P* = 1.3×10^-7^) and *diseases of esophagus* (*P* = 4.3×10^-7^)*. Muscle weakness* (*P =* 2.8×10^-6^) and *abnormal movements* (*P =* 3.9×10^-6^) are consistent with neurological traits reported in 16p11.2 deletion carriers such as hypotonia and motor impairments ^81^. *Sleep apnea* (*P =* 8.9×10^-6^) was a novel phenotype, potentially related to increased BMI in deletion carriers. 16p11.2 duplication carrier status was similarly associated with developmental diagnoses (Figure 3): *lack of normal physiological development* (*P* = 5.6×10^-15^), *developmental delays and disorders* (*P* = 2.5×10^-13^), *delayed milestones* (*P* = 9.0×10^-13^), *autism* (*P* = 1.3×10^-12^), and *mental retardation* (*P* = 1.6×10^-7^) ^3,5^. 16p11.2 duplication carriers status was also associated with multiple heart defects, including *valvular heart disease/heart chambers* (*P =* 4.6×10^-10^) and *cardiac shunt/heart septal defect* (*P* = 3.2×10^-8^), both of which have been reported previously ^82^. 16p11.2 duplications are known to be a risk factor for epilepsy, and were associated with an epilepsy-related diagnosis of *convulsions* (*P* = 2.9×10^-8^) in the biobank ^3,83^. *Infantile cerebral palsy* (*P* = 4.9×10^-6^), while a potential reason for genetic testing, has not previously been associated with 16p11.2 duplications. While the 16p11.2 CNV contains genes such as *SPN* and *MVP* that are active in the immune system, there is no prior evidence of the susceptibility of duplication carriers to infection, making the diagnosis *Bacterial infection NOS* (*P* = 5.5×10^-7^) a novel finding.

For 22q11.2 deletion carriers, the canonical associated features were cardiac defects such as *cardiomegaly* (*P* = 3.5×10^-258^) and *cardiac shunt/heart septal defects* (*P* = 4.7×10^-285^) (Figure 4) ^6,7^. Other highly associated diagnoses were developmental: *lack of normal physiological development* (*P* = 1.7×10^-47^)*, developmental delays and disorders* (*P =* 6.3×10^-29^)*, delayed milestones* (*P* = 6.0×10^-11^) ^6,7^. Congenital anomalies such as *cleft palate* (*P =* 9.4×10^-80^) were also over-represented. The secondary known traits for 22q11.2 deletion carriers included *immunity deficiency* (*P <* 10^-285^), and *disorders involving the immune mechanism* (*P* < 10^-285^). Previously, it has been reported that 50% of 22q11.2 deletion carriers have T-cell dysfunction and 17% have humoral dysfunction ^7^. Very few traits over-represented in 22q11.2 deletion carriers were novel; one of these was *hyperpotassemia* (*P* = 1.4×10^-10^).

22q11.2 duplication carrier status was also associated with developmental diagnoses (Figure 4): *delayed milestones* (*P* = 1.1×10^-13^), l*ack of normal physiological development* (*P* = 9.7×10^-13^)*, pervasive developmental disorders* (*P* = 1.2×10^-6^) ^8^. 22q11.2 duplication status was associated with cardiac phenotypes such as *Cardiac shunt/ heart septal defect* (*P =* 2.3×10^-5^). Cardiac features have not as often been reported in 22q11.2 duplication carriers compared to 22q11.2 deletion carriers ^8^. Remaining traits such as *abnormality of gait* (*P =* 3.1×10^-12^) and *hearing loss* (*P =* 2.1×10^-7^) have also been seen in 22q11.2, including as indications for genetic testing ^84^.

### Phenome-wide association studies identify phenotypic consequences of expression variation in 16p11.2 and 22q11.2 genes

As our study of the impact of the entire CNV on phenotype confirmed our ability to detect important CNV-associated traits within the BioVU biobank, our next goal was to catalogue how each individual CNV gene might affect the medical phenome. We generated predicted expression for CNV and flanking genes, as in the initial GWAS analyses, for the 48,630 non-CNV-carrier individuals genotyped in BioVU. We tested 1,531 medical phenotypic codes meeting frequency criteria (n = 20 cases) in this subset. There were six phenome-wide significant (*P* < 3.3×10^-5^) gene-trait associations at 16p11.2 including: *INO80E* with *skull and face fracture and other intercranial injury* (*P* = 1.9×10^-15^), *NPIPB11* with *psychosis* (*P* = 1.0×10^-5^), and *SLX1B* with *psychosis* (*P* = 3.0×10^-5^). There were eleven phenome-wide significant gene-trait associations at 22q11.2 including: *AIFM3* with *renal failure (P =* 2.3×10^-5^), *LZTR1* with *malignant neoplasm, other* (*P =* 1.4×10^-5^), *SCARF2* with *mood disorders* (*P =* 1.3×10^-5^), *PI4KA* with *disorders of iris and ciliary body* (*P* = 1.1×10^-7^) and *disorders resulting from impaired renal function* (*P* = 2.2×10^-5^). These include two renal traits, consistent with the 22q11.2 deletion carrier status association with *renal failure.* The associations of *LZTR1* and *PI4KA* with neoplasms and eye disorders correspond to similar traits associated with these genes in prior literature ^108–110^.

Previously established gene-trait associations came up as suggestive (top 1 percentile), though not phenome-wide significant, associations in this cohort. *TBX1*, a gene at 22q11.2 tied to heart development, had *other chronic ischemic heart disease, unspecified* (*P =* 0.001), *endocarditis* (*P =* 0.0046), *cardiomyopathy* (*P =* 0.0055), and *coronary atherosclerosis* (*P =* 0.0076) among its top 1% phenome associations ^29–32^. *TBX6* at 16p11.2, which has a role in bone development and scoliosis, has *pathologic fracture of vertebrae* in its top 1% phenome associations (*P =* 0.0028) ^85–87^. *TANGO2* mutations at 22q11.2 have been associated with metabolic abnormalities such as hypoglycemia, and our PheWAS for *TANGO2* showed *abnormal glucose* (*P =* 0.0013) as a top phenotype ^88, 90^.

As few gene-trait pairs reached phenome-wide significance and established associations were present at more nominal levels, we also considered traits that did not meet the significance threshold in our analysis, but were in the top 1% of phenotypic associations for a given gene (Table S8). We found that traits categorized as “mental disorders” were over-represented in the top 1% of the phenome of CNV genes (*P* = 5.2×10^-5^). Of all 17 clinical categories tested, “mental disorders” was the only trait with enrichment p-value meeting multiple testing thresholds (Table S9). This suggested that the effect of CNV genes is more widespread on brain-related traits than simply those detected as statistically significant.

Some of the top 1% PheWAS traits for CNV genes overlapped with the original five traits we studied: schizophrenia, IQ, BMI, bipolar disorder, and ASD. At 16p11.2 (Table 1, Figure 5), there were genes whose top PheWAS results included schizophrenia-related traits (*Psychosis, Schizophrenia and other psychotic disorders),* IQ-related traits (*Developmental delays and disorders, Mental retardation, Delayed milestones*), BMI-related traits (*Bariatric surgery, Morbid obesity*), and ASD-related traits (*Pervasive developmental disorders*). At 22q11.2 (Table 2, Figure 5), there were genes whose top PheWAS results included schizophrenia-related traits (*Hallucinations*), BMI-related traits (*Overweight, obesity and other hyperalimentation, Morbid obesity*), ASD-related traits (*Autism, Speech and language disorder*), and bipolar-related traits (*Mood disorders*). We could not perform strict independent replication for these associations because many of these traits are difficult to define in the same way across datasets (for example *speech and language disorder* vs. *autism*). Instead, we compared the top association statistics within our GWAS discovery and replication datasets for the genes identified to be associated with brain-related traits in PheWAS as an extension of this study (Table S5). The following genes were associated at *P* < 0.05 and also in the top 5^th^ percentile within at least one of the GWAS discovery or replication datasets (Table S5): *SEPT1* (*psychosis* – in UK Biobank schizophrenia 20002_1289 *P* = 0.03), *AIFM3* (*mood disorders* – in UK Biobank bipolar F31 *P =* 0.04), *SCARF2* (*mood disorders* – in UK Biobank bipolar F31 *P* = 0.003), *HIC2* (*mood disorders* – in UK Biobank bipolar 20002_1991, *P =* 0.004), *ZNF48* (*bariatric surgery* – in UK Biobank BMI 3.7×10^-6^). Of these, the association between *SCARF2* and *mood disorders* reached phenome-wide significance in the PheWAS.

**Figure 5.**
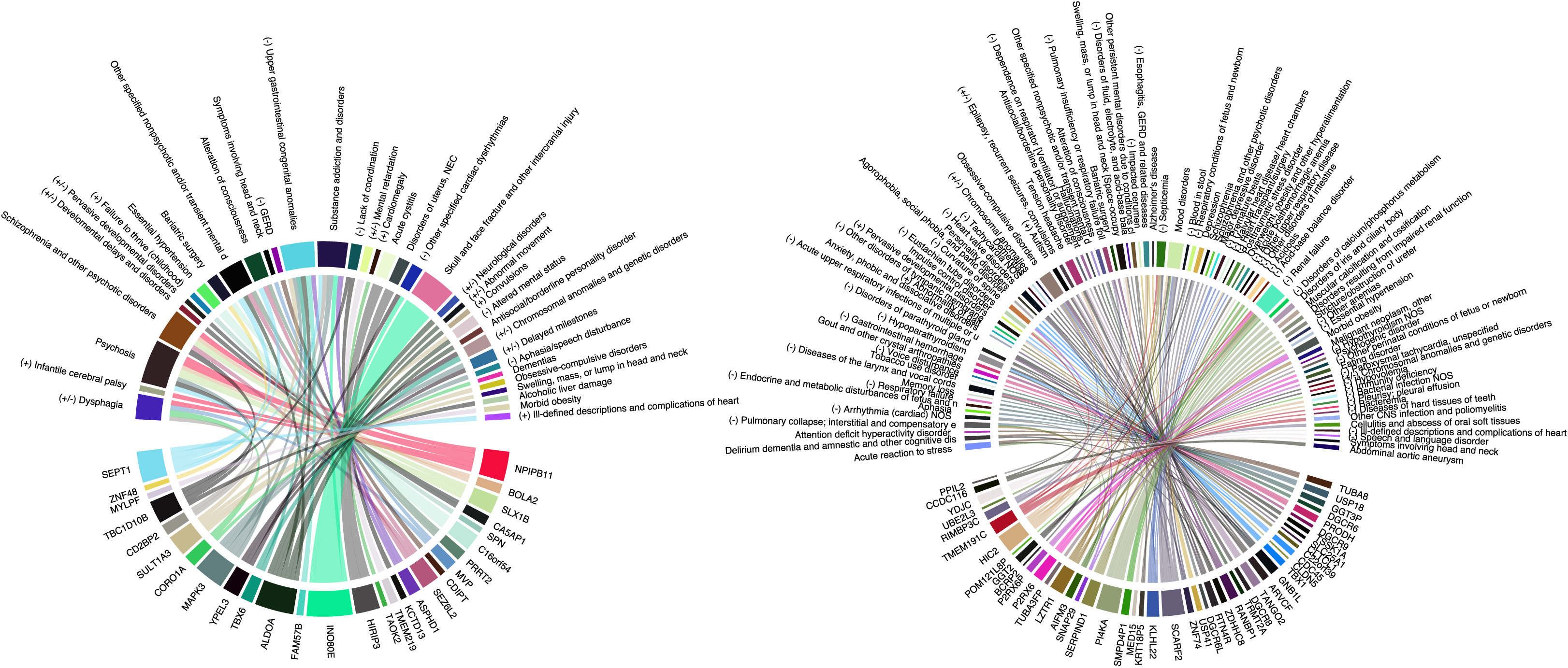
Graphical summary of selected PheWAS results by gene. Each circle contains the CNV genes, in chromosomal order, on the bottom, and their associated PheWAS traits at the top. Genes are connected to their PheWAS-associated traits, with the width of the line proportional to the -log10 p-value of the association. If a trait is also over-represented in duplication and/or deletion carriers it is marked with a + (duplications), - (deletions), or +/- (both). The complete list of gene-trait pairs can be found in Tables 1 and 2, and Supplemental Table S7.

Predicted expressions may be correlated between nearby genes, thus multiple genes can share a PheWAS trait association due to correlation alone. We are underpowered for independence testing for the majority of our GWAS traits, but we selected several notable traits that appeared in multiple genes to test for independence, in the same way as in our GWAS analysis (Table S5). We performed a conditional analysis on 16p11.2 genes whose top phenome associations included *psychosis: NPIPB11, BOLA2, MAPK3, SEPT1, SLX1B, TBC1D10B.* By comparing whether the p-value of association stayed constant vs. increased after conditioning, we found that *NPIPB11, SEPT1, SLX1B, TBC1D10B* were likely independent associations, whereas *BOLA2* and *MAPK3* may be associated with *psychosis* at least partly by correlation with the other four. We also performed the same analysis for 22q11.2 genes whose top phenome associations included *morbid obesity*: *SNAP29, P2RX6, P2RX6P.* Of these genes, the only one with a p-value increase was *P2RX6P*, suggesting that its association with *morbid obesity* may be explained at least in part by another gene. From conditional analysis, we see evidence of a multigenic contribution to both of these traits from CNV genes.

### Genes in 16p11.2 and 22q11.2 are associated with traits that are also over-represented in carriers

We originally hypothesized that small variations in CNV gene expression would be associated with phenotypes similar to those that were present in CNV carriers, perhaps with smaller effects. Our use of electronic health records first on the entire CNV itself, then on individual genes allows us to detect these potential effects across traits. Unlike the five brain-related traits that we originally chose, many of the traits in the EHRs do not have similar large GWAS datasets available. Considering that our non-ascertained biobank is not well-powered for less common traits, we chose to focus on the top one percentile of the phenome associations rather than the few associations that passed the phenome-wide significance threshold.

Traits that were found both in 16p11.2 carriers and in individual genes’ PheWAS results included primary CNV traits such as *mental retardation* and *delayed milestones*, as well as secondary traits such as *dysphagia* and *convulsions* (Table 1, Figure 5). There were six genes (*ASPHD1, FAM57B, ALDOA, TBX6, MAPK3, SULT1A3)* whose top PheWAS associations included the 16p11.2 deletion-associated trait of *upper gastrointestinal congenital anomalies*, though we are underpowered to know whether all of these signals are independent. Of the genes that we found as drivers in the first analysis of GWAS datasets, we note that *INO80E*’s top PheWAS results overlap the 16p11.2 deletion-associated trait *other specified cardiac dysrhythmias* and *SPN*’s top PheWAS results overlap the 16p11.2 duplication-associated trait of *failure to thrive (childhood*).

Over 30 genes at 22q11.2 had a top PheWAS trait overlapping a trait over-represented in 22q11.2 duplication or deletion carriers (Table 2, Figure 5). Top PheWAS results for 22q11.2 genes included primary cardiac traits such as *tachycardia* (*P2RX6P, GNB1L*) and primary brain-related traits such as *autism* (*TANGO2, ZDHHC8).* We also found genes with top PheWAS results overlapping secondary traits from the carrier screen, such as *diseases of the larynx and vocal cords* (*DGCR6, PRODH, ARVCF*).

We also tested whether the top associations from individual gene PheWAS results were enriched for EHR phenotypes over-represented in carriers. We did this by analyzing where top PheWAS traits associated with CNV genes were ranked within PheWAS results of carrier status. We found no evidence for enrichment in 16p11.2 duplications, 16p11.2 deletions, 22q11.2 duplications, or 22q11.2 deletions (Figure S4).

## DISCUSSION

In this study, we sought to identify individual genes in the 16p11.2 and 22q11.2 regions driving brain-related disorders, as well as the impact of both the entire CNV and specific CNV genes on the medical phenome. In a novel *in-silico* approach to CNV fine-mapping, we tested whether genetically-driven predicted expression variation of the individual genes in each CNV was associated with brain-related disorders ascertained in GWAS data. We identified individual genes at 16p11.2 whose expression was associated with schizophrenia (*INO80E*), IQ (*SPN*), and BMI (*SPN, INO80E*) in the expected direction based on known 16p11.2 biology. We then used EHR data to detect (known and novel) traits overrepresented in 16p11.2 and 22q11.2 carriers for comparison with individual gene results. Third, we used the same EHR system biobank containing over 1,500 medical traits to explore the consequences of expression variation of 16p11.2 and 22q11.2 CNV genes in non-carriers, and identified enrichment of brain-related traits as well as individual genes potentially driving carrier-associated traits. The results from the GWAS-derived and PheWAS analyses can be considered as two independent ways to probe the function of CNV genes using expression imputation.

*INO80E,* the gene we identified as a driver of schizophrenia and BMI, is a chromatin remodeling gene and has rarely been considered in the context of brain-related traits ^94^. Mice heterozygous for this gene have shown abnormal locomotor activation ^95^. Locomotor activity in mice is a frequently used proxy for brain-related disorders including schizophrenia ^96^. Our results are consistent with a previous observation that eQTLs from dorsolateral prefrontal cortex for *INO80E* co-localize with schizophrenia GWAS SNPs ^97^. In addition, an analogous imputed expression based transcriptome-wide association study observed association between *INO80E* and schizophrenia using summary statistics ^98^. A third transcriptomic association study using prenatal and adult brain tissues also pointed to *INO80E* as a risk gene for schizophrenia ^99^. By focusing on a specific schizophrenia-associated region, using individual level data, and performing a conditional analysis, we have obtained additional precision, and were able to fine-map the signal at 16p11.2 down to a single gene. Our study differs from Gusev *et al* and Walker *et al* in the expression prediction models used: we used 48 tissue models from the Genotype-Tissue Expression consortium, Gusev *et al* used brain, blood, and adipose tissues from other consortia, and Walker *et al* used prenatal and adult brain tissues only. The overlap in association results shows that our approach is robust to variation in predictive models. Furthermore, we find that the utilization of non-brain tissues in our analysis did not hinder our ability to detect this association. Mice with a heterozygous mutation in *Ino80e* showed increased body weight, consistent with our BMI association result for the same gene ^95^.

*SPN*, a gene highly associated with both IQ and BMI, is active in immune cells and is not known to play a role in brain-related disorders ^100, 101^. We note that the association p-values for *SPN* are much lower than for any other genes showing association signal. This may be because our approach detected relatively few eQTLs for *SPN* (12 SNPs in two tissues), many of which overlapped with highly associated GWAS SNPs for both IQ and BMI, rather than contributing to noise.

Our results give evidence that pleiotropy is involved in the pathogenicity of 16p11.2, as opposed to a strictly “one gene, one trait” model. Specifically, *INO80E* was associated with both schizophrenia and BMI, and *SPN* was associated with both BMI and IQ. Genetic correlations of at least -0.05 and as much as -0.5 have been estimated for the BMI/IQ and SCZ/BMI pairs, suggesting that pleiotropy may play a general role in these disorders ^102–105^. Consistent with the genetic correlations, most (8/12) eQTL SNPs in our prediction models for *SPN* drove the associations with both IQ and BMI.

While most associations we detected were in the expected direction given previous knowledge, *MVP* and *KCTD13* were associated with BMI in the opposite (positive) direction, and *YPEL3* with schizophrenia in the negative direction. We resolved the schizophrenia result by conditional analysis, where we found that *YPEL3* was associated with schizophrenia simply due to correlation with *INO80E*. For BMI, we were able to use UK Biobank data to determine that *MVP* was not an independent association with BMI, while *KCTD13* remained. For an example like *KCTD13,* we offer three explanations: these results may be false-positives due to correlation-based “hitchhiking”, they may demonstrate a limitation of our approach, or they may have a true BMI-increasing effect. First, we cannot rule out that it “hitchhikes” to statistical significance with other negatively-associated genes due to correlation, but does not contribute to BMI itself. Second, this result might represent a limitation of our eQTL-based method. *KCTD13* is a highly brain-expressed gene, but had no high-quality brain prediction models ^51^. The direction of the eQTLs regulating *KCTD13* expression in the brain may be brain-specific, and brain may be the only relevant tissue for the effect of *KCTD13* on BMI. That is, *KCTD13* may actually have a strong negative correlation with BMI, but falsely appears positive due to the specific eQTLs used for expression prediction. Such tissue-specific eQTL directions of effect have been observed for at least 2,000 genes ^106^. Improved brain-specific prediction models will resolve this limitation. Third, *KCTD13* could have a true BMI-increasing effect. If so, the 16p11.2 region contains both BMI-increasing and BMI-decreasing genes, and the effect of the BMI-decreasing genes is stronger. Such a model is a potential explanation for the observation that duplications at 16p11.2 in mice, unlike humans, are associated with obesity ^44^. One set of genes may be the more influential determinant of the obesity trait in each organism.

Our PheWAS of traits overrepresented in 16p11.2 and 22q11.2 carriers served as a validation of our biobank EHR approach via detection of previously identified CNV-associated traits. Brain-related traits, such as *delayed milestones, mental retardation* and *pervasive developmental disorders*, were among the top over-represented traits in both 16p11.2 and 22q11.2 CNV carriers. 22q11.2 deletion carriers were strongly associated with *cardiac congenital anomalies* and *cleft palate*, two of the hallmark features of the CNV. Even though the total number of CNV carriers within the biobank was relatively small, the overrepresentation was strong enough to let us pick up these differences. At the same time, we identified novel traits that may be confirmed in larger samples of CNV carriers such as *sleep apnea* in 16p11.2 deletions and *hyperpotassemia* in 22q11.2 deletions.

Our PheWAS between the predicted expressions of 16p11.2 and 22q11.2 genes and 1,500 medical phenotypic codes resulted in 17 phenome-wide significant gene-trait pairs. Some of these genes have been shown to drive similar traits in prior literature. The gene *AIFM3* at 22q11.2 was associated with *renal failure. AIFM3* is a gene in a proposed critical region for 22q11.2-associated kidney defects, and led to kidney defects in zebrafish ^107^. *SNAP29,* another gene associated with kidney defects in the same study, had *renal failure, NOS* in its top 1% phenome associations. *LZTR1* was significantly associated with *malignant neoplasm, other.* This gene is a cause of schwannomatosis, a disease involving neoplasms (albeit normally benign) ^108^. Model organisms with defects in *PI4KA,* associated with *disorders of iris and ciliary body* in our study, showed eye-related phenotypes ^109, 110^. Because few genes had any associations which were phenome-wide significant, we elected to analyze the top 1% of associations of each gene. We noticed that our gene-by-gene PheWAS recapitulated the effect of *TBX1* on the circulatory system and of *TBX6* on the musculoskeletal system at this threshold ^29–31, 85–87^. We have not found any evidence that *TBX1* contributes to canonical 22q11.2-associated brain-related traits. Notably, we found that clinical traits in the *mental disorders* category were over-represented in the top 1% of associations among all genes tested, and *mental disorders* was the only category significantly enriched. Some mental disorders, such as *psychosis*, were top PheWAS hits for multiple genes, but we were underpowered for rigorous independence testing. Still, we have identified potential genetic drivers for each of those traits. Moreover, three novel brain-related gene-trait pairs reached phenome-wide significance: *NPIPB11* and *SLX1B* near the CNV breakpoint at 16p11.2 with *psychosis*, as well as *SCARF2* at 22q11.2 with *mood disorders.* The expression of *SLX1B* is modified in 16p11.2 carriers; *NPIPB11* expression differences have not been detected in transcriptomic studies of 16p11.2 ^44, 46^. Integrating genetic information with the diagnosis of *mood disorders* in the clinical data allowed us to find a new candidate, *SCARF2*, at 22q11.2 that we were unable or underpowered to detect in the ascertained bipolar data alone.

### Limitations

The 16p11.2 and 22q11.2 CNVs are significant risk factors for ASD and schizophrenia, respectively, and yet no individual genes in either CNV were associated with case-control status for the associated trait in the best-powered datasets available to us. Assuming the true causal gene(s) for these disorders do exist within the CNV, limitations in our approach may preclude us from discovering them. Predictive models for gene expression are imperfect; while they capture the cis-heritability of gene expression, they may not capture the entire variability of the expression of a gene (the largest single-tissue prediction R^2^ for our genes is 0.45, and the average R^2^ is 0.07). Moreover, there are genes in both regions for which no high-quality models exist. If the causal gene is among the genes that cannot be well predicted, we cannot detect this gene by our approach. One category of genes that are not represented in our study are microRNAs. 22q11.2 carriers have a unique microRNA signature, and the contribution of microRNA to 22q11.2-CNV associated schizophrenia has been previously hypothesized ^111, 112^. If the microRNAs are important regulatory elements for 22q11.2-associated traits, our approach is insufficient to detect them.

Rather than focusing on any specific tissue(s), we chose to perform a cross-tissue analysis, an approach that improves power to detect gene-trait associations and detected 16p11.2 genes associated with schizophrenia, IQ, and BMI ^59^. While we might expect that brain-specific models would be best at detecting relevant genes for brain-related traits, we are limited by the amount of data available – brain tissue transcriptomes are available for fewer than half of the GTEx individuals ^52^. An underlying assumption behind the use of all tissues (rather than just brain)tissues for these mental disorders is that eQTLs for our genes of interest are shared across tissues, and that the same eQTLs affect the expression of a gene in the brain as in other tissues. In general, eQTLs tend to be either highly shared between tissues or highly tissue-specific, largely as a function of the gene being expressed exclusively or nearly exclusively in a single tissue^113^. The GTEx correlation of eQTL effect sizes between brain and non-brain tissues is 0.499 (Spearman) ^52^. We may miss genes of interest that have brain-specific expression but not enough power to detect eQTLs. Furthermore, as these eQTLs come from adult tissues, we would miss genes where effects on brain-related traits are specific to early developmental timepoints.

A further limitation is that the variation in expression that can be modeled using eQTLs may be considerably smaller for some genes than the effect of deletions and duplications. For example, there may be a gene at 22q11.2 for which decreases in expression contribute to schizophrenia, but only when expression levels are reduced to nearly 50% of the expression levels of non-carriers. Furthermore, the expression predictions of these genes are calculated solely using *cis*- eQTLs within 1MB of the gene ^58^. It may be necessary to consider the effect of *trans*-eQTLs to explore the genetic effect of expression variation accurately. Similarly, we have not considered *trans-*effects due to chromosome contacts, such as those that exist between the 16p11.2 region described here and another smaller CNV region elsewhere at 16p11.2 ^114, 115^.

Because the CNV carrier individuals in our biobank are young (median age < 18), we don’t yet know what traits might commonly occur once individuals reach older age. There were traits in our analysis that were over-represented in older carriers, but difficult to interpret as they didn’t meet our frequency threshold, including: *dementia with cerebral degenerations* in 22q11.2 deletion carriers, *anterior horn cell disease* in 16p11.2 deletion carriers, and *cerebral degenerations, unspecified* in 16p11.2 duplication carriers. These findings show a need for longitudinal studies of carrier cohorts and studies of carriers in older age. Such additional data may point to additional clinical features of 16p11.2 and 22q11.2 CNV carriers.

So far, we have considered the effect of each CNV gene independently, when, in reality, the genes may not be acting independently. A *Drosophila* model for 16p11.2 genes has shown evidence of epistasis between genes within a CNV as a modifier of phenotype ^40^. If there are 16p11.2 traits in humans also driven by epistasis, our single-gene screen would not have detected the appropriate genes for those traits.

### Conclusions

In developing our approach, we hypothesized that naturally occurring variation in gene expression of CNV genes in non-carriers would convey risk for traits seen in CNV carriers. We found that this was true for at least three 16p11.2 associated traits: BMI, schizophrenia, and IQ. Promisingly, the direction of association was generally consistent with whether the trait was found in duplication or deletion carriers. Our approach is computationally efficient, extendable to other CNV-trait pairs, and overcomes one limitation of animal models by testing the effect of CNV genes specifically in humans.

In this study, we synthesized information from both large GWAS studies and EHR-linked biobanks, benefiting from the strengths of both approaches. Psychiatric brain-related disorders such as autism, schizophrenia, and bipolar disorder have a population frequency below 5%, so large datasets specifically ascertained for brain-related disorders are better at providing sufficient statistical power for association analysis, especially when the effect of each gene is small. On the other hand, the presence of many diagnostic codes in a biobank help identify brain-related traits that may be relevant to CNVs but not the primary reported symptoms, such as speech and language disorder. We were also able to carry out two distinct and complementary analyses using the same dataset. The presence of CNV carrier status in the EHR-linked biobank allowed us to probe the phenotypic consequences of the entire deletion or duplication. Then, we were able to test each CNV gene for association with the same diagnostic descriptions.

Our novel approach provided insights into how individual genes in the 16p11.2 and 22q11.2 CNVs may drive health and behavior in a human population. Expression imputation methods allowed us to study the predicted effects of individual CNV genes in large human populations. The incorporation of medical records into biobanks provided a way to determine clinical symptoms and diagnoses to which expression differences in the genes may contribute. We expect our ability to detect genes with this type of approach to increase in the coming years, as more individuals in biobanks are genotyped, the number of individuals contributing to large cohorts grow, and the methods to more finely and accurately predict gene expression improve. Additional experiments on our newly prioritized genes are necessary to determine their specific functional impact on brain-related disorders and to evaluate their value as putative therapeutic targets.

## SUPPLEMENTAL DATA

As supplement to this manuscript, we include four figures, nine tables, and one document listing consortia members.

## ACKNOWLEDGEMENTS

We would like to acknowledge the consortia and individuals that provided the individual data we used in this study: BioVU at Vanderbilt University, the Genotype-Tissue Expression Consortium, and the Autism, Schizophrenia, and Bipolar working groups of the Psychiatric Genomics Consortium. PGC working group authors are listed in Note S1. We would also like to thank Megan Sheuy for providing cleaned BMI data for BioVU participants. This work was sponsored by the National Institute of Mental Health [R01 MH107467 to LAW, R01 MH113362 to NJC]; as well as the National Human Genome Research Institute [U01 HG009086 to NJC]. TWM was also supported by the National Human Genome Research Institute [T32 HG008341]. The funders had no role in study design, data collection and analysis, decision to publish, or preparation of the manuscript. The BioVU projects at Vanderbilt University Medical Center are supported by numerous sources: institutional funding, private agencies, and federal grants. These include the NIH funded Shared Instrumentation Grant S10OD017985 and S10RR025141; CTSA grants UL1TR002243, UL1TR000445, and UL1RR024975 from the National Center for Advancing Translational Sciences. Its contents are solely the responsibility of the authors and do not necessarily represent official views of the National Center for Advancing Translational Sciences or the National Institutes of Health. Genomic data are also supported by investigator-led projects that include U01HG004798, R01NS032830, RC2GM092618, P50GM115305, U01HG006378, U19HL065962, R01HD074711; and additional funding sources listed at https://victr.vumc.org/biovu-funding/.

## DATA AVAILABILITY

Individual-level genotypes for Psychiatric Genomics Consortium cohorts can be obtained by submitting an application at www.med.unc.edu/pgc/. Summary level genetic datasets used here are available to freely download from GIANT BMI (https://portals.broadinstitute.org/collaboration/giant/index.php/GIANT_consortium) and CNCR IQ (https://ctg.cncr.nl/software/summary_statistics). BioVU phenotypes are available by application through Vanderbilt University. GTEx genotypes and phenotypes are listed on dbGAP (phs000424.v7.p2). The complete results of our analyses of these datasets are available in the supplement.

---

Figure S1: Position of the 16p11.2 and 22q11.2 CNVs on chromosomes 16 and 22 and list of genes in these regions. Bolded genes had predictive models available and were considered in our study. 22q11.2 genes are staggered on either side of the chromosome for visual clarity.

Figure S2: Association between 16p11.2 genes and the remaining two brain-related traits. Association between predicted expression of 16p11.2 genes and ASD (left), bipolar disorder (right). Genes are listed on the horizontal axis in order of chromosomal position. The -log10 p-values on the vertical axis are given a positive or negative direction based on the average direction of the single-tissue results. The significance threshold, *P* < 7.6×10^-5^, is a Bonferroni correction on the total number of 16p11.2 and 22q11.2 genes (132) tested across 5 traits (0.05/(5*132)).

Figure S3: Association between 22q11.2 genes and five brain-related traits. Association between predicted expression of 22q11.2 genes and, from bottom to top, ASD, bipolar disorder, schizophrenia, BMI, and IQ, using MultiXcan (ASD, bipolar disorder, schizophrenia) and S-MultiXcan (BMI, IQ). Genes are listed on the horizontal axis in order of chromosomal position. The -log10 p-values on the vertical axis are given a positive or negative direction based on the average direction of the single-tissue results. The significance threshold, *P* < 7.6×10^-5^, is a Bonferroni correction on the total number of 16p11.2 and 22q11.2 genes (132) tested across 5 traits (0.05/(5*132)).

Figure S4: Test of enrichment of gene-based PheWAS results within carrier vs. non-carrier PheWAS results. The histograms show the distribution of ranks of the traits associated with individual 16p11.2 or 22q11.2 genes within the ranks of traits from the association analysis with carrier status. A Kolmogorov-Smirnov test was applied to determine whether the distribution of ranks of traits associated with individual CNV genes was different from the expected (i.e. uniform) distribution of ranks of traits associated with carrier status. The x-axis shows counts of traits from the gene-by-gene PheWAS. and the y-axis shows ranks. The gene-based PheWAS results would be considered enriched within carrier status PheWAS results if their ranks tended to skew towards one side of the distribution.

Note S1: Supplemental Acknowledgements. Members of the Psychiatric Genomics Consortium who contributed to this work.

Table S1: List of genotyped discovery and replication cohorts used in the study.

List of datasets used for discovery and replication of association results with sample sizes. The specific cohorts from the Psychiatric Genomics Consortium that were used for this analysis are listed. All variables from the UK Biobank that were used for replication are shown.

Table S2: List of genes at or near 16p11.2 and 22q11.2.

List of coding and non-coding genes in the CNV region, as well as flanking genes 200kb on either side. Genes for which PrediXcan models based on GTEx v7 were available and the range of model qualities (R^2^) are noted, along with the number of tissues in which prediction models were available. Genes are annotated with their type (e.g. protein-coding, pseudogene, etc.), whether they are in the CNV or flanking, and any other names by which they may be referred in the literature.

Table S3: Identifying 16p11.2 and 22q11.2 cases from electronic health records (EHR). Keyword searches across all documents within the Vanderbilt EHR were performed to identify individuals carrying 16p11.2 or 22q11.2 CNVs. Individuals with documents containing matching keywords were reviewed manually to confirm the presence of 16p11.2 or 22q11.2 CNV. Individuals were excluded from case groups if their records included a mention of additional CNVs. Individuals within the 16p11.2 case groups were also excluded if the size of the reported CNV was 200-250 kb. Individuals within the 22q11.2 case group were excluded if the size of the CNV was smaller than 500 kb or if there was a mention of “distal” when referring to the deletion or duplication. Confirmed case numbers are listed, with the non-genotyped counts in parentheses. Non-genotyped individuals were used for downstream phenome-wide analyses.

Table S4: Results of MultiXcan and S-MultiXcan associations between CNV genes and autism, schizophrenia, bipolar disorder, BMI, and IQ.

For autism, bipolar disorder, and schizophrenia, z-scores and p-values come from a METAL meta-analysis across PGC cohorts. For BMI and IQ, mean z-scores and p-values come directly from S-MultiXcan output. Genes in each CNV are sorted by chromosomal position.

Table S5: Conditional analysis for independence of associations.

Conditional analysis was performed on the PGC schizophrenia data, the UK Biobank BMI data, as well as two BioVU clinical trait associations (16p11.2 genes and *psychosis*, 22q11.2 genes and *morbid obesity*). For each trait, we performed MultiXcan first adjusting for a specific gene, then by leaving a gene in and adjusting all of the other genes associated with that trait out. The *P_cond_* reported in the text is the p-value of this gene-trait pair when adjusting for all other genes considered for conditioning for this trait, unless otherwise stated.

Table S6: Comparison of association results to independent data.

For each gene-trait pair, we list the original p-value, the GWAS trait(s) that we classified as most similar to a PheWAS trait, its best p-value in an independent dataset, the number of GWAS datasets that were used for this trait, and the rank of this gene within that dataset. For UK Biobank summary statistics, we have genome-wide data; for datasets with individual-level data, only 16p11.2 and 22q11.2 genes were calculated. See Table S2 for more information on datasets used.

Table S7: Traits over-represented in CNV carriers.

The four categories of CNV carrier – 16p11.2 duplication, 16p11.2 deletion, 22q11.2 duplication, 22q11.2 deletion – were tested separately. The results for all clinical traits tested are provided. The number of cases and controls for each trait is given, as well as whether the p-value meets either Bonferroni or FDR correction. Traits in bold were represented in over 5% of carriers.

Table S8: Top PheWAS associations of 16p11.2 and 22q11.2 genes.

The top 15 associated traits for each gene, regardless of p-value, are shown. These represent the top 1% of associations among all traits tested. Genes are listed in alphabetical order, with each trait’s sample size and phecode (www.phewascatalog.org) noted.

Table S9: Enrichment of clinical categories among the top PheWAS associations.

The top 15 traits (codes) for each gene analyzed (n = 1470 gene-trait pairs) were divided into 17 clinical categories (observed counts column). The values in the expected counts column are calculated as 1470 * {the proportion of traits of that category tested}. For example, 159 out of 1531 codes tested were from the “circulatory system” category, so the expected counts for “circulatory system” are calculated as 1470*159/1531. The last column contains the p-value from a binomial test comparing whether the observed proportion of clinical categories is more extreme than expected.

## REFERENCES

1. Jacquemont, S., Reymond, A., Zufferey, F., Harewood, L., Walters, R.G., Kutalik, Z., Martinet, D., Shen, Y., Valsesia, A., Beckmann, N.D., et al. (2011). Mirror extreme BMI phenotypes associated with gene dosage at the chromosome 16p11.2 locus. Nature 478, 97–102.

2. McCarthy, S.E., Makarov, V., Kirov, G., Addington, A.M., McClellan, J., Yoon, S., Perkins, D.O., Dickel, D.E., Kusenda, M., Krastoshevsky, O., et al. (2009). Microduplications of 16p11.2 are associated with schizophrenia. Nat. Genet. 41, 1223–1227.

3. Shinawi, M., Liu, P., Kang, S.-H.L., Shen, J., Belmont, J.W., Scott, D.A., Probst, F.J., Craigen, W.J., Graham, B.H., Pursley, A., et al. (2010). Recurrent reciprocal 16p11.2 rearrangements associated with global developmental delay, behavioural problems, dysmorphism, epilepsy, and abnormal head size. J. Med. Genet. 47, 332–341.

4. Kumar, R.A., KaraMohamed, S., Sudi, J., Conrad, D.F., Brune, C., Badner, J.A., Gilliam, T.C., Nowak, N.J., Cook, E.H., Dobyns, W.B., et al. (2007). Recurrent 16p11.2 microdeletions in autism. Hum. Mol. Genet. 17, 628–638.

5. Weiss, L.A., Shen, Y., Korn, J.M., Arking, D.E., Miller, D.T., Fossdal, R., Saemundsen, E., Stefansson, H., Ferreira, M.A.R., Green, T., et al. (2008). Association between Microdeletion and Microduplication at 16p11.2 and Autism. N. Engl. J. Med. 358, 667–675.

6. Bassett, A.S., Chow, E.W.C., Husted, J., Weksberg, R., Caluseriu, O., Webb, G.D., and Gatzoulis, M.A. (2005). Clinical features of 78 adults with 22q11 deletion syndrome. Am. J. Med. Genet. Part A 138A, 307–313.

7. Campbell, I.M., Sheppard, S.E., Crowley, T.B., McGinn, D.E., Bailey, A., McGinn, M.J., Unolt, M., Homans, J.F., Chen, E.Y., Salmons, H.I., et al. (2018). What is new with 22q? An update from the 22q and You Center at the Children’s Hospital of Philadelphia. Am. J. Med. Genet. Part A 176, 2058–2069.

8. Wentzel, C., Fernström, M., Öhrner, Y., Annerén, G., and Thuresson, A.-C. (2008). Clinical variability of the 22q11.2 duplication syndrome. Eur. J. Med. Genet. 51, 501–510.

9. Schneider, M., Debbané, M., Bassett, A.S., Chow, E.W.C., Fung, W.L.A., van den Bree, M.B.M., Owen, M., Murphy, K.C., Niarchou, M., Kates, W.R., et al. (2014). Psychiatric Disorders From Childhood to Adulthood in 22q11.2 Deletion Syndrome: Results From the International Consortium on Brain and Behavior in 22q11.2 Deletion Syndrome. Am. J. Psychiatry 171, 627–639.

10. Voll, S.L., Boot, E., Butcher, N.J., Cooper, S., Heung, T., Chow, E.W.C., Silversides, C.K., and Bassett, A.S. (2017). Obesity in adults with 22q11.2 deletion syndrome. Genet. Med. 19, 204–208.

11. Carlson, C., Papolos, D., Pandita, R.K., Faedda, G.L., Veit, S., Goldberg, R., Shprintzen, R., Kucherlapati, R., and Morrow, B. (1997). Molecular analysis of velo-cardio-facial syndrome patients with psychiatric disorders. Am. J. Hum. Genet. 60, 851–859.

12. Sahoo, T., Theisen, A., Rosenfeld, J.A., Lamb, A.N., Ravnan, J.B., Schultz, R.A., Torchia, B.S., Neill, N., Casci, I., Bejjani, B.A., et al. (2011). Copy number variants of schizophrenia susceptibility loci are associated with a spectrum of speech and developmental delays and behavior problems. Genet. Med. 13, 868–880.

13. Itsara, A., Cooper, G.M., Baker, C., Girirajan, S., Li, J., Absher, D., Krauss, R.M., Myers, R.M., Ridker, P.M., Chasman, D.I., et al. (2008). Population analysis of large copy number variants and hotspots of human genetic disease. Am. J. Hum. Genet. 84, 148–161.

14. Bijlsma, E.K., Gijsbers, A.C.J., Schuurs-Hoeijmakers, J.H.M., van Haeringen, A., Fransen van de Putte, D.E., Anderlid, B.-M., Lundin, J., Lapunzina, P., Pérez Jurado, L.A., Delle Chiaie, B., et al. (2009). Extending the phenotype of recurrent rearrangements of 16p11.2: Deletions in mentally retarded patients without autism and in normal individuals. Eur. J. Med. Genet. 52, 77– 87.

15. McCarthy, S.E., Makarov, V., Kirov, G., Addington, A.M., McClellan, J., Yoon, S., Perkins, D.O., Dickel, D.E., Kusenda, M., Krastoshevsky, O., et al. (2009). Microduplications of 16p11.2 are associated with schizophrenia. Nat. Genet. 41, 1223–1227.

16. Rees, E., Kirov, G., Sanders, A., Walters, J.T.R., Chambert, K.D., Shi, J., Szatkiewicz, J., O’Dushlaine, C., Richards, A.L., Green, E.K., et al. (2014). Evidence that duplications of 22q11.2 protect against schizophrenia. Mol. Psychiatry 19, 37–40.

17. Walters, R.G., Jacquemont, S., Valsesia, A., de Smith, A.J., Martinet, D., Andersson, J., Falchi, M., Chen, F., Andrieux, J., Lobbens, S., et al. (2010). A new highly penetrant form of obesity due to deletions on chromosome 16p11.2. Nature 463, 671–675.

18. Smith, A.C.M., McGavran, L., Robinson, J., Waldstein, G., Macfarlane, J., Zonona, J., Reiss, J., Lahr, M., Allen, L., Magenis, E., et al. (1986). Interstitial deletion of (17)(p11.2p11.2) in nine patients. Am. J. Med. Genet. 24, 393–414.

19. Potocki, L., Chen, K.S., Park, S.S., Osterholm, D.E., Withers, M.A., Kimonis, V., Summers, A.M., Meschino, W.S., Anyane-Yeboa, K., Kashork, C.D., et al. (2000). Molecular mechanism for duplication 17p11.2 - The homologous recombination reciprocal of the Smith-Magenis microdeletion. Nat. Genet. 24, 84–87.

20. Slager, R.E., Newton, T.L., Vlangos, C.N., Finucane, B., and Elsea, S.H. (2003). Mutations in RAI1 associated with Smith–Magenis syndrome. Nat. Genet. 33, 466–468.

21. Walz, K., Paylor, R., Yan, J., Bi, W., and Lupski, J.R. (2006). Rai1 duplication causes physical and behavioral phenotypes in a mouse model of dup(17)(p11.2p11.2). J. Clin. Invest. 116, 3035–3041.

22. Williams, J.C.P., Barratt-Boyes, B.G., and Lowe, J.B. (1961). Supravalvular Aortic Stenosis. Circulation 24, 1311–1318.

23. Beuren, A.J., Apitz, J., and Harmjanz, D. (1962). Supravalvular Aortic Stenosis in Association with Mental Retardation and a Certain Facial Appearance. Circulation 26, 1235– 1240.

24. Curran, M.E., Atkinson, D.L., Ewart, A.K., Morris, C.A., Leppert, M.F., and Keating, M.T. (1993). The elastin gene is disrupted by a translocation associated with supravalvular aortic stenosis. Cell 73, 159–168.

25. Ewart, A.K., Morris, C.A., Atkinson, D., Jin, W., Sternes, K., Spallone, P., Stock, A.D., Leppert, M., and Keating, M.T. (1993). Hemizygosity at the elastin locus in a developmental disorder, Williams syndrome. Nat. Genet. 5, 11–16.

26. Koolen, D.A., Vissers, L.E.L.M., Pfundt, R., De Leeuw, N., Knight, S.J.L., Regan, R., Kooy, R.F., Reyniers, E., Romano, C., Fichera, M., et al. (2006). A new chromosome 17q21.31 microdeletion syndrome associated with a common inversion polymorphism. Nat. Genet. 38, 999–1001.

27. Koolen, D.A., Sharp, A.J., Hurst, J.A., Firth, H. V, Knight, S.J.L., Goldenberg, A., Saugier-Veber, P., Pfundt, R., Vissers, L.E.L.M., Destrée, A., et al. (2008). Clinical and molecular delineation of the 17q21.31 microdeletion syndrome. J. Med. Genet. 45, 710–720.

28. Koolen, D.A., Kramer, J.M., Neveling, K., Nillesen, W.M., Moore-Barton, H.L., Elmslie, F. V, Toutain, A., Amiel, J., Malan, V., Tsai, A.C.-H., et al. (2012). Mutations in the chromatin modifier gene KANSL1 cause the 17q21.31 microdeletion syndrome. Nat. Genet. 44, 639–641.

29. Jerome, L.A., and Papaioannou, V.E. (2001). DiGeorge syndrome phenotype in mice mutant for the T-box gene, Tbx1. Nat. Genet. 27, 286–291.

30. Lindsay, E.A., Vitelli, F., Su, H., Morishima, M., Huynh, T., Pramparo, T., Jurecic, V., Ogunrinu, G., Sutherland, H.F., Scambler, P.J., et al. (2001). Tbx1 haploinsufficiency in the DiGeorge syndrome region causes aortic arch defects in mice. Nature 410, 97–101.

31. Merscher, S., Funke, B., Epstein, J.A., Heyer, J., Puech, A., Lu, M.M., Xavier, R.J., Demay, M.B., Russell, R.G., Factor, S., et al. (2001). TBX1 Is Responsible for Cardiovascular Defects in Velo-Cardio-Facial/DiGeorge Syndrome. Cell 104, 619–629.

32. Paylor, R., Glaser, B., Mupo, A., Ataliotis, P., Spencer, C., Sobotka, A., Sparks, C., Choi, C.- H., Oghalai, J., Curran, S., et al. (2006). Tbx1 haploinsufficiency is linked to behavioral disorders in mice and humans: implications for 22q11 deletion syndrome. Proc. Natl. Acad. Sci. U. S. A. 103, 7729–7734.

33. Ma, G., Shi, Y., Tang, W., He, Z., Huang, K., Li, Z., He, G., Feng, G., Li, H., and He, L. (2007). An association study between the genetic polymorphisms within TBX1 and schizophrenia in the Chinese population.

34. Consortium, S.W.G. of the P.G., Ripke, S., Walters, J.T., and O’Donovan, M.C. (2020). Mapping genomic loci prioritises genes and implicates synaptic biology in schizophrenia. MedRxiv 2020.09.12.20192922.

35. Clements, C.C., Wenger, T.L., Zoltowski, A.R., Bertollo, J.R., Miller, J.S., de Marchena, A.B., Mitteer, L.M., Carey, J.C., Yerys, B.E., Zackai, E.H., et al. (2017). Critical region within 22q11.2 linked to higher rate of autism spectrum disorder. Mol. Autism 8, 58.

36. Crepel, A., Steyaert, J., De la Marche, W., De Wolf, V., Fryns, J.-P., Noens, I., Devriendt, K., and Peeters, H. (2011). Narrowing the critical deletion region for autism spectrum disorders on 16p11.2. Am. J. Med. Genet. Part B Neuropsychiatr. Genet. 156, 243–245.

37. Pucilowska, J., Vithayathil, J., Tavares, E.J., Kelly, C., Colleen Karlo, J., and Landreth, G.E. (2015). The 16p11.2 deletion mouse model of autism exhibits altered cortical progenitor proliferation and brain cytoarchitecture linked to the ERK MAPK pathway. J. Neurosci. 35, 3190–3200.

38. Golzio, C., Willer, J., Talkowski, M.E., Oh, E.C., Taniguchi, Y., Jacquemont, S., Reymond, A., Sun, M., Sawa, A., Gusella, J.F., et al. (2012). KCTD13 is a major driver of mirrored neuroanatomical phenotypes of the 16p11.2 copy number variant. Nature 485, 363–367.

39. Blaker-Lee, A., Gupta, S., McCammon, J.M., De Rienzo, G., and Sive, H. (2012). Zebrafish homologs of genes within 16p11.2, a genomic region associated with brain disorders, are active during brain development, and include two deletion dosage sensor genes. Dis. Model. Mech. 5,.

40. Iyer, J., Singh, M.D., Jensen, M., Patel, P., Pizzo, L., Huber, E., Koerselman, H., Weiner, A.T., Lepanto, P., Vadodaria, K., et al. (2018). Pervasive genetic interactions modulate neurodevelopmental defects of the autism-Associated 16p11.2 deletion in Drosophila melanogaster. Nat. Commun. 9, 1–19.

41. Paylor, R., McIlwain, K.L., McAninch, R., Nellis, A., Yuva-Paylor, L.A., Baldini, A., and Lindsay, E.A. (2001). Mice deleted for the DiGeorge/velocardiofacial syndrome region show abnormal sensorimotor gating and learning and memory impairments. Hum. Mol. Genet. 10, 2645–2650.

42. Guna, A., Butcher, N.J., and Bassett, A.S. (2015). Comparative mapping of the 22q11.2 deletion region and the potential of simple model organisms. J. Neurodev. Disord. 7, 18.

43. McCammon, J.M., Blaker-Lee, A., Chen, X., and Sive, H. (2017). The 16p11.2 homologs fam57ba and doc2a generate certain brain and body phenotypes. Hum. Mol. Genet. 26, 3699– 3712.

44. Arbogast, T., Ouagazzal, A.-M., Chevalier, C., Kopanitsa, M., Afinowi, N., Migliavacca, E., Cowling, B.S., Birling, M.-C., Champy, M.-F., Reymond, A., et al. (2016). Reciprocal Effects on Neurocognitive and Metabolic Phenotypes in Mouse Models of 16p11.2 Deletion and Duplication Syndromes. PLOS Genet. 12, e1005709.

45. Ward, T.R., Zhang, X., Leung, L.C., Zhou, B., Muench, K., Roth, J.G., Khechaduri, A., Plastini, M.J., Charlton, C., Pattni, R., et al. (2020). Genome-wide molecular effects of the neuropsychiatric 16p11 CNVs in an iPSC-to-iN neuronal model. BioRxiv 2020.02.09.940965.

46. Blumenthal, I., Ragavendran, A., Erdin, S., Klei, L., Sugathan, A., Guide, J.R., Manavalan, P., Zhou, J.Q., Wheeler, V.C., Levin, J.Z., et al. (2014). Transcriptional Consequences of 16p11.2 Deletion and Duplication in Mouse Cortex and Multiplex Autism Families. Am. J. Hum. Genet. 94, 870–883.

47. Luo, R., Sanders, S.J., Tian, Y., Voineagu, I., Huang, N., Chu, S.H., Klei, L., Cai, C., Ou, J., Lowe, J.K., et al. (2012). Genome-wide Transcriptome Profiling Reveals the Functional Impact of Rare De Novo and Recurrent CNVs in Autism Spectrum Disorders. Am. J. Hum. Genet. 91, 38–55.

48. Zhang, X., Zhang, Y., Zhu, X., Purmann, C., Haney, M.S., Ward, T., Khechaduri, A., Yao, J., Weissman, S.M., and Urban, A.E. (2018). Local and global chromatin interactions are altered by large genomic deletions associated with human brain development. Nat. Commun. 9, 5356.

49. Jalbrzikowski, M., Lazaro, M.T., Gao, F., Huang, A., Chow, C., Geschwind, D.H., Coppola, G., and Bearden, C.E. (2015). Transcriptome Profiling of Peripheral Blood in 22q11.2 Deletion Syndrome Reveals Functional Pathways Related to Psychosis and Autism Spectrum Disorder. PLoS One 10, e0132542.

50. Migliavacca, E., Golzio, C., Männik, K., Blumenthal, I., Oh, E.C., Harewood, L., Kosmicki, J.A., Loviglio, M.N., Giannuzzi, G., Hippolyte, L., et al. (2015). A Potential Contributory Role for Ciliary Dysfunction in the 16p11.2 600 kb BP4-BP5 Pathology. Am. J. Hum. Genet. 96, 784– 796.

51. Aguet, F., Barbeira, A.N., Bonazzola, R., Brown, A., Castel, S.E., Jo, B., Kasela, S., Kim-Hellmuth, S., Liang, Y., Oliva, M., et al. (2019). The GTEx Consortium atlas of genetic regulatory effects across human tissues. BioRxiv 787903.

52. Aguet, F., Ardlie, K.G., Cummings, B.B., Gelfand, E.T., Getz, G., Hadley, K., Handsaker, R.E., Huang, K.H., Kashin, S., Karczewski, K.J., et al. (2017). Genetic effects on gene expression across human tissues. Nature 550, 204–213.

53. Lonsdale, J., Thomas, J., Salvatore, M., Phillips, R., Lo, E., Shad, S., Hasz, R., Walters, G., Garcia, F., Young, N., et al. (2013). The Genotype-Tissue Expression (GTEx) project. Nat. Genet. 45, 580–585.

54. Ardlie, K.G., DeLuca, D.S., Segrè, A. V., Sullivan, T.J., Young, T.R., Gelfand, E.T., Trowbridge, C.A., Maller, J.B., Tukiainen, T., Lek, M., et al. (2015). The Genotype-Tissue Expression (GTEx) pilot analysis: Multitissue gene regulation in humans. Science (80-.). 348, 648–660.

55. Freund, M.K., Burch, K., Shi, H., Mancuso, N., Kichaev, G., Garske, K.M., Pan, D.Z., Pajukanta, P., Pasaniuc, B., and Arboleda, V.A. Phenotype-specific enrichment of Mendelian disorder genes near GWAS regions across 62 complex traits.

56. Lupski, J.R., Belmont, J.W., Boerwinkle, E., and Gibbs, R.A. (2011). Clan genomics and the complex architecture of human disease. Cell 147, 32–43.

57. Blair, D.R., Lyttle, C.S., Mortensen, J.M., Bearden, C.F., Jensen, A.B., Khiabanian, H., Melamed, R., Rabadan, R., Bernstam, E. V., Brunak, S., et al. (2013). A nondegenerate code of deleterious variants in mendelian loci contributes to complex disease risk. Cell 155, 70–80.

58. Gamazon, E.R., Wheeler, H.E., Shah, K.P., Mozaffari, S. V., Aquino-Michaels, K., Carroll, R.J., Eyler, A.E., Denny, J.C., Nicolae, D.L., Cox, N.J., et al. (2015). A gene-based association method for mapping traits using reference transcriptome data. Nat. Genet. 47, 1091–1098.

59. Barbeira, A.N., Pividori, M.D., Zheng, J., Wheeler, H.E., Nicolae, D.L., and Im, H.K. (2019). Integrating predicted transcriptome from multiple tissues improves association detection. PLOS Genet. 15, e1007889.

60. Schizophrenia Working Group of the Psychiatric Genomics Consortium, S. (2014). Biological insights from 108 schizophrenia-associated genetic loci. Nature 511, 421–427.

61. Stahl, E.A., Breen, G., Forstner, A.J., McQuillin, A., Ripke, S., Trubetskoy, V., Mattheisen, M., Wang, Y., Coleman, J.R.I., Gaspar, H.A., et al. (2019). Genome-wide association study identifies 30 loci associated with bipolar disorder. Nat. Genet. 51, 793–803.

62. Grove, J., Ripke, S., Als, T.D., Mattheisen, M., Walters, R.K., Won, H., Pallesen, J., Agerbo, E., Andreassen, O.A., Anney, R., et al. (2019). Identification of common genetic risk variants for autism spectrum disorder. Nat. Genet. 51, 431–444.

63. Locke, A.E., Kahali, B., Berndt, S.I., Justice, A.E., Pers, T.H., Day, F.R., Powell, C., Vedantam, S., Buchkovich, M.L., Yang, J., et al. (2015). Genetic studies of body mass index yield new insights for obesity biology. Nature 518, 197–206.

64. Savage, J.E., Jansen, P.R., Stringer, S., Watanabe, K., Bryois, J., De Leeuw, C.A., Nagel, M., Awasthi, S., Barr, P.B., Coleman, J.R.I., et al. (2018). Genome-wide association meta-analysis in 269,867 individuals identifies new genetic and functional links to intelligence. Nat. Genet. 50, 912–919.

65. Roden, D., Pulley, J., Basford, M., Bernard, G., Clayton, E., Balser, J., and Masys, D. (2008). Development of a Large-Scale De-Identified DNA Biobank to Enable Personalized Medicine. Clin. Pharmacol. Ther. 84, 362–369.

66. Schizophrenia Working Group of the Psychiatric Genomics Consortium, {fname} (2014). Biological insights from 108 schizophrenia-associated genetic loci. Nature 511, 421–427.

67. Yengo, L., Sidorenko, J., Kemper, K.E., Zheng, Z., Wood, A.R., Weedon, M.N., Frayling, T.M., Hirschhorn, J., Yang, J., and Visscher, P.M. (2018). Meta-analysis of genome-wide association studies for height and body mass index in ∼700 000 individuals of European ancestry. Hum. Mol. Genet. 27, 3641–3649.

68. Barbeira, A.N., Dickinson, S.P., Bonazzola, R., Zheng, J., Wheeler, H.E., Torres, J.M., Torstenson, E.S., Shah, K.P., Garcia, T., Edwards, T.L., et al. (2018). Exploring the phenotypic consequences of tissue specific gene expression variation inferred from GWAS summary statistics. Nat. Commun. 9, 1825.

69. Aguet, F., Brown, A.A., Castel, S.E., Davis, J.R., He, Y., Jo, B., Mohammadi, P., Park, Y.S., Parsana, P., Segrè, A. V., et al. (2017). Genetic effects on gene expression across human tissues. Nature 550, 204–213.

70. Willer, C.J., Li, Y., and Abecasis, G.R. (2010). METAL: fast and efficient meta-analysis of genomewide association scans. Bioinformatics 26, 2190–2191.

71. Roden, D.M., Pulley, J.M., Basford, M.A., Bernard, G.R., Clayton, E.W., Balser, J.R., and Masys, D.R. (2008). Development of a large-scale de-identified DNA biobank to enable personalized medicine. Clin. Pharmacol. Ther. 84, 362–369.

72. McCarthy, S., Das, S., Kretzschmar, W., Delaneau, O., Wood, A.R., Teumer, A., Kang, H.M., Fuchsberger, C., Danecek, P., Sharp, K., et al. (2016). A reference panel of 64,976 haplotypes for genotype imputation. Nat. Genet. 48, 1279–1283.

73. Das, S., Forer, L., Schönherr, S., Sidore, C., Locke, A.E., Kwong, A., Vrieze, S.I., Chew, E.Y., Levy, S., McGue, M., et al. (2016). Next-generation genotype imputation service and methods. Nat. Genet. 48, 1284–1287.

74. Denny, J.C., Ritchie, M.D., Basford, M.A., Pulley, J.M., Bastarache, L., Brown-Gentry, K., Wang, D., Masys, D.R., Roden, D.M., and Crawford, D.C. (2010). PheWAS: demonstrating the feasibility of a phenome-wide scan to discover gene-disease associations. Bioinformatics 26, 1205–1210.

75. Carroll, R.J., Bastarache, L., and Denny, J.C. (2014). R PheWAS: data analysis and plotting tools for phenome-wide association studies in the R environment. Bioinformatics 30, 2375–2376.

76. Gu, Z., Gu, L., Eils, R., Schlesner, M., and Brors, B. (2014). circlize implements and enhances circular visualization in R. Bioinformatics 30, 2811–2812.

77. Dantas, A.G., Santoro, M.L., Nunes, N., de Mello, C.B., Pimenta, L.S.E., Meloni, V.A., Soares, D.C.Q., Belangero, S.N., Carvalheira, G., Kim, C.A., et al. (2019). Downregulation of genes outside the deleted region in individuals with 22q11.2 deletion syndrome. Hum. Genet. 138, 93–103.

78. UK Biobank — Neale lab.

79. Denny, J.C., Bastarache, L., Ritchie, M.D., Carroll, R.J., Zink, R., Mosley, J.D., Field, J.R., Pulley, J.M., Ramirez, A.H., Bowton, E., et al. (2013). Systematic comparison of phenome-wide association study of electronic medical record data and genome-wide association study data. Nat. Biotechnol. 31, 1102–1111.

80. Roth, J.G., Muench, K.L., Asokan, A., Mallett, V.M., Gai, H., Verma, Y., Weber, S., Charlton, C., Fowler, J.L., Loh, K.M., et al. (2020). Copy Number Variation at 16p11.2 Imparts Transcriptional Alterations in Neural Development in an hiPSC-derived Model of Corticogenesis. BioRxiv 2020.04.22.055731.

81. Steinman, K.J., Spence, S.J., Ramocki, M.B., Proud, M.B., Kessler, S.K., Marco, E.J., Green Snyder, L., D’Angelo, D., Chen, Q., Chung, W.K., et al. (2016). 16p11.2 deletion and duplication: Characterizing neurologic phenotypes in a large clinically ascertained cohort. Am. J. Med. Genet. Part A 170, 2943–2955.

82. Karunanithi, Z., Vestergaard, E.M., and Lauridsen, M.H. (2017). Transposition of the great arteries - a phenotype associated with 16p11.2 duplications? World J. Cardiol. 9, 848–852.

83. Fernandez, B.A., Roberts, W., Chung, B., Weksberg, R., Meyn, S., Szatmari, P., Joseph-George, A.M., MacKay, S., Whitten, K., Noble, B., et al. (2010). Phenotypic spectrum associated with de novo and inherited deletions and duplications at 16p11.2 in individuals ascertained for diagnosis of autism spectrum disorder. J. Med. Genet. 47, 195–203.

84. Wenger, T.L., Miller, J.S., DePolo, L.M., de Marchena, A.B., Clements, C.C., Emanuel, B.S., Zackai, E.H., McDonald-McGinn, D.M., and Schultz, R.T. (2016). 22q11.2 duplication syndrome: elevated rate of autism spectrum disorder and need for medical screening. Mol. Autism 7, 27.

85. Watabe-Rudolph, M., Schlautmann, N., Papaioannou, V.E., and Gossler, A. (2002). The mouse rib-vertebrae mutation is a hypomorphic Tbx6 allele. Mech. Dev. 119, 251–256.

86. Chen, W., Liu, J., Yuan, D., Zuo, Y., Liu, Z., Liu, S., Zhu, Q., Qiu, G., Huang, S., Giampietro, P.F., et al. (2016). Progress and perspective of TBX6 gene in congenital vertebral malformations. Oncotarget 7, 57430–57441.

87. Liu, J., Wu, N., Yang, N., Takeda, K., Chen, W., Li, W., Du, R., Liu, S., Zhou, Y., Zhang, L., et al. (2019). TBX6-associated congenital scoliosis (TACS) as a clinically distinguishable subtype of congenital scoliosis: further evidence supporting the compound inheritance and TBX6 gene dosage model. Genet. Med. 21, 1548–1558.

88. Lalani, S.R., Liu, P., Rosenfeld, J.A., Watkin, L.B., Chiang, T., Leduc, M.S., Zhu, W., Ding, Y., Pan, S., Vetrini, F., et al. (2016). Recurrent Muscle Weakness with Rhabdomyolysis, Metabolic Crises, and Cardiac Arrhythmia Due to Bi-allelic TANGO2 Mutations. Am. J. Hum. Genet. 98, 347–357.

89. Li, Y., Wu, Z., Xu, L., Feng, Z., Wang, Y., Dai, Z., Liu, Z., Sun, X., Qiu, Y., and Zhu, Z. (2020). Genetic Variant of TBX1 Gene is Functionally Associated with Adolescent Idiopathic Scoliosis in the Chinese Population. Spine (Phila. Pa. 1976).

90. Dines, J.N., Golden-Grant, K., LaCroix, A., Muir, A.M., Cintrón, D.L., McWalter, K., Cho, M.T., Sun, A., Merritt, J.L., Thies, J., et al. (2019). TANGO2: expanding the clinical phenotype and spectrum of pathogenic variants. Genet. Med. 21, 601–607.

91. Bittel, D.C., Yu, S., Newkirk, H., Kibiryeva, N., Holt 3rd, A., Butler, M.G., and Cooley, L.D. (2009). Refining the 22q11.2 deletion breakpoints in DiGeorge syndrome by aCGH. Cytogenet. Genome Res. 124, 113–120.

92. Thomas, R.A., Ambalavanan, A., Rouleau, G.A., and Barker, P.A. (2016). Identification of genetic variants of LGI1 and RTN4R (NgR1) linked to schizophrenia that are defective in NgR1-LGI1 signaling. Mol. Genet. Genomic Med. 4, 447–456.

93. Kou, I., Otomo, N., Takeda, K., Momozawa, Y., Lu, H.F., Kubo, M., Kamatani, Y., Ogura, Y., Takahashi, Y., Nakajima, M., et al. (2019). Genome-wide association study identifies 14 previously unreported susceptibility loci for adolescent idiopathic scoliosis in Japanese. Nat. Commun. 10, 1–9.

94. Ayala, R., Willhoft, O., Aramayo, R.J., Wilkinson, M., McCormack, E.A., Ocloo, L., Wigley, D.B., and Zhang, X. (2018). Structure and regulation of the human INO80-nucleosome complex. Nature 556, 391–395.

95. Bult, C.J., Blake, J.A., Smith, C.L., Kadin, J.A., and Richardson, J.E. (2018). Mouse Genome Database (MGD) 2019. Nucleic Acids Res. 47, D801–D806.

96. Powell, C.M., and Miyakawa, T. (2006). Schizophrenia-relevant behavioral testing in rodent models: a uniquely human disorder? Biol. Psychiatry 59, 1198–1207.

97. Dobbyn, A., Huckins, L.M., Boocock, J., Sloofman, L.G., Glicksberg, B.S., Giambartolomei, C., Hoffman, G.E., Perumal, T.M., Girdhar, K., Jiang, Y., et al. (2018). Landscape of Conditional eQTL in Dorsolateral Prefrontal Cortex and Co-localization with Schizophrenia GWAS. Am. J. Hum. Genet. 102, 1169–1184.

98. Gusev, A., Mancuso, N., Won, H., Kousi, M., Finucane, H.K., Reshef, Y., Song, L., Safi, A., McCarroll, S., Neale, B.M., et al. (2018). Transcriptome-wide association study of schizophrenia and chromatin activity yields mechanistic disease insights. Nat. Genet. 50, 538–548.

99. Walker, R.L., Ramaswami, G., Hartl, C., Mancuso, N., Gandal, M.J., de la Torre-Ubieta, L., Pasaniuc, B., Stein, J.L., and Geschwind, D.H. (2019). Genetic Control of Expression and Splicing in Developing Human Brain Informs Disease Mechanisms. Cell 179, 750–771.e22.

100. Pallant, A., Eskenazi, A., Mattei, M.G., Fournier, R.E.K., Carlsson, S.R., Fukuda, M., and Frelinger, J.G. (1989). Characterization of cDNAs encoding human leukosialin and localization of the leukosialin gene to chromosome 16. Proc. Natl. Acad. Sci. U. S. A. 86, 1328–1332.

101. Park, J.K., Rosenstein, Y.J., Remold-O’Donnell, E., Bierer, B.E., Rosen, F.S., and Burakoff, S.J. (1991). Enhancement of T-cell activation by the CD43 molecule whose expression is defective in Wiskott-Aldrich syndrome. Nature 350, 706–709.

102. Marioni, R.E., Yang, J., Dykiert, D., Mõttus, R., Campbell, A., Davies, G., Hayward, C., Porteous, D.J., Visscher, P.M., and Deary, I.J. (2016). Assessing the genetic overlap between BMI and cognitive function. Mol. Psychiatry 21, 1477–1482.

103. Sabia, S., Kivimaki, M., Shipley, M.J., Marmot, M.G., and Singh-Manoux, A. (2009). Body mass index over the adult life course and cognition in late midlife: the Whitehall II Cohort Study. Am. J. Clin. Nutr. 89, 601–607.

104. Ikeda, M., Tanaka, S., Saito, T., Ozaki, N., Kamatani, Y., and Iwata, N. (2018). Re-evaluating classical body type theories: Genetic correlation between psychiatric disorders and body mass index. Psychol. Med. 48, 1745–1748.

105. Bulik-Sullivan, B., Finucane, H.K., Anttila, V., Gusev, A., Day, F.R., Loh, P.R., Duncan, L., Perry, J.R.B., Patterson, N., Robinson, E.B., et al. (2015). An atlas of genetic correlations across human diseases and traits. Nat. Genet. 47, 1236–1241.

106. Mizuno, A., and Okada, Y. (2019). Biological characterization of expression quantitative trait loci (eQTLs) showing tissue-specific opposite directional effects. Eur. J. Hum. Genet.

107. Lopez-Rivera, E., Liu, Y.P., Verbitsky, M., Anderson, B.R., Capone, V.P., Otto, E.A., Yan, Z., Mitrotti, A., Martino, J., Steers, N.J., et al. (2017). Genetic Drivers of Kidney Defects in the DiGeorge Syndrome. N. Engl. J. Med. NEJMoa1609009.

108. Piotrowski, A., Xie, J., Liu, Y.F., Poplawski, A.B., Gomes, A.R., Madanecki, P., Fu, C., Crowley, M.R., Crossman, D.K., Armstrong, L., et al. (2014). Germline loss-of-function mutations in LZTR1 predispose to an inherited disorder of multiple schwannomas. Nat. Genet. 46, 182–187.

109. Ma, H., Blake, T., Chitnis, A., Liu, P., and Balla, T. (2009). Crucial role of phosphatidylinositol 4-kinase IIIα in development of zebrafish pectoral fin is linked to phosphoinositide 3-kinase and FGF signaling. J. Cell Sci. 122, 4303–4310.

110. Bojjireddy, N., Botyanszki, J., Hammond, G., Creech, D., Peterson, R., Kemp, D.C., Snead, M., Brown, R., Morrison, A., Wilson, S., et al. (2014). Pharmacological and genetic targeting of the PI4KA enzyme reveals its important role in maintaining plasma membrane phosphatidylinositol 4-phosphate and phosphatidylinositol 4,5-bisphosphate levels. J. Biol. Chem. 289, 6120–6132.

111. Forstner, A.J., Degenhardt, F., Schratt, G., and Nöthen, M.M. (2013). MicroRNAs as the cause of schizophrenia in 22q11.2 deletion carriers, and possible implications for idiopathic disease: a mini-review. Front. Mol. Neurosci. 6, 47.

112. De la Morena, M.T., Eitson, J.L., Dozmorov, I.M., Belkaya, S., Hoover, A.R., Anguiano, E., Pascual, M.V., and van Oers, N.S.C. (2013). Signature MicroRNA expression patterns identified in humans with 22q11.2 deletion/DiGeorge syndrome. Clin. Immunol. 147, 11–22.

113. Consortium, T.Gte. (2020). The GTEx Consortium atlas of genetic regulatory effects across human tissues. Science 369, 1318–1330.

114. Loviglio, M.N., Leleu, M., Männik, K., Passeggeri, M., Giannuzzi, G., van der Werf, I., Waszak, S.M., Zazhytska, M., Roberts-Caldeira, I., Gheldof, N., et al. (2016). Chromosomal contacts connect loci associated with autism, BMI and head circumference phenotypes. Mol. Psychiatry.

115. Bachmann-Gagescu, R., Mefford, H.C., Cowan, C., Glew, G.M., Hing, A. V, Wallace, S., Bader, P.I., Hamati, A., Reitnauer, P.J., Smith, R., et al. (2010). Recurrent 200-kb deletions of 16p11.2 that include the SH2B1 gene are associated with developmental delay and obesity. Genet. Med. 12, 641–647.

