## Supplementary figures and images for "Integration of genetic, transcriptomic, and clinical data provides insight into 16p11.2 and 22q11.2 CNV genes"

### Table_S3.docx

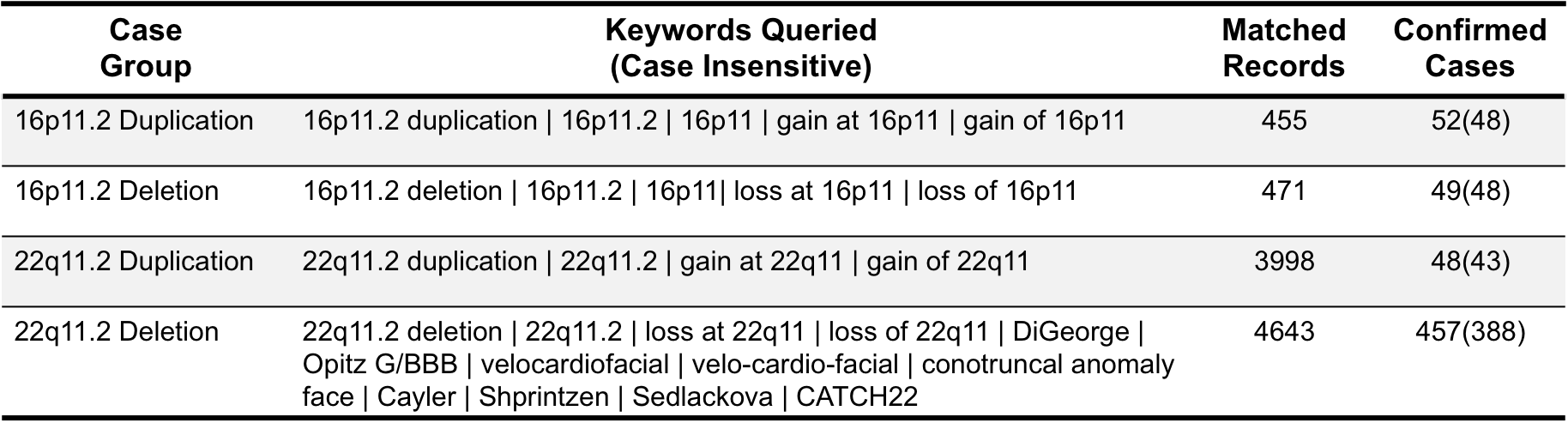
