## Supplemental Figures and Note for "Integration of genetic, transcriptomic, and clinical data provides insight into 16p11.2 and 22q11.2 CNV genes"

**Legends to supplemental figures and tables**

Figure S1: Position of the 16p11.2 and 22q11.2 CNVs on chromosomes 16 and 22 and list of genes in these regions. Bolded genes had predictive models available and were considered in our study. 22q11.2 genes are staggered on either side of the chromosome for visual clarity.

Figure S2: Association between 16p11.2 genes and the remaining two brain-related traits. Association between predicted expression of 16p11.2 genes and ASD (left), bipolar disorder (right). Genes are listed on the horizontal axis in order of chromosomal position. The -log10 p-values on the vertical axis are given a positive or negative direction based on the average direction of the single-tissue results. The significance threshold, $P < 7.6 \times 10^{-5}$, is a Bonferroni correction on the total number of 16p11.2 and 22q11.2 genes (132) tested across 5 traits (0.05/(5*132)).

Figure S3: Association between 22q11.2 genes and five brain-related traits.
Association between predicted expression of 22q11.2 genes and, from bottom to top, ASD, bipolar disorder, schizophrenia, BMI, and IQ, using MultiXcan (ASD, bipolar disorder, schizophrenia) and S-MultiXcan (BMI, IQ). Genes are listed on the horizontal axis in order of chromosomal position. The -log10 p-values on the vertical axis are given a positive or negative direction based on the average direction of the single-tissue results. The significance threshold, $P < 7.6 \times 10^{-5}$, is a Bonferroni correction on the total number of 16p11.2 and 22q11.2 genes (132) tested across 5 traits (0.05/(5*132)).

Figure S4: Test of enrichment of gene-based PheWAS results within carrier vs. non-carrier PheWAS results. The histograms show the distribution of ranks of the traits associated with individual 16p11.2 or 22q11.2 genes within the ranks of traits from the association analysis with carrier status. A Kolmogorov-Smirnov test was applied to determine whether the distribution of ranks of traits associated with individual CNV genes was different from the expected (i.e. uniform) distribution of ranks of traits associated with carrier status. The x-axis shows counts of traits from the gene-by-gene PheWAS. and the y-axis shows ranks. The gene-based PheWAS results would be considered enriched within carrier status PheWAS results if their ranks tended to skew towards one side of the distribution.

Note S1: Supplementary Acknowledgements. Members of the Psychiatric Genomics Consortium who contributed to this work.

Table S1: List of genotyped discovery and replication cohorts used in the study.
List of datasets used for discovery and replication of association results with sample sizes. The specific cohorts from the Psychiatric Genomics Consortium that were used for this analysis are listed. All variables from the UK Biobank that were used for replication are shown.

Table S2: List of genes at or near 16p11.2 and 22q11.2.
List of coding and non-coding genes in the CNV region, as well as flanking genes 200kb on either side. Genes for which PrediXcan models based on GTEx v7 were available and the range of model qualities ($R^2$) are noted, along with the number of tissues in which prediction models

were available. Genes are annotated with their type (e.g. protein-coding, pseudogene, etc.), whether they are in the CNV or flanking, and any other names by which they may be referred in the literature.

Table S3: Identifying 16p11.2 and 22q11.2 cases from electronic health records (EHR**).** Keyword searches across all documents within the Vanderbilt EHR were performed to identify individuals carrying 16p11.2 or 22q11.2 CNVs. Individuals with documents containing matching keywords were reviewed manually to confirm the presence of 16p11.2 or 22q11.2 CNV. Individuals were excluded from case groups if their records included a mention of additional CNVs. Individuals within the 16p11.2 case groups were also excluded if the size of the reported CNV was 200-250 kb. Individuals within the 22q11.2 case group were excluded if the size of the CNV was smaller than 500 kb or if there was a mention of "distal" when referring to the deletion or duplication. Confirmed case numbers are listed, with the non-genotyped counts in parentheses. Non-genotyped individuals were used for downstream phenome-wide analyses.

Table S4: Results of MultiXcan and S-MultiXcan associations between CNV genes and autism, schizophrenia, bipolar disorder, BMI, and IQ.

For autism, bipolar disorder, and schizophrenia, z-scores and p-values come from a METAL meta-analysis across PGC cohorts. For BMI and IQ, mean z-scores and p-values come directly from S-MultiXcan output. Genes in each CNV are sorted by chromosomal position.

Table S5: Conditional analysis for independence of associations.

Conditional analysis was performed on the PGC schizophrenia data, the UK Biobank BMI data, as well as two BioVU clinical trait associations (16p11.2 genes and *psychosis*, 22q11.2 genes and *morbid obesity*). For each trait, we performed MultiXcan first adjusting for a specific gene, then by leaving a gene in and adjusting all of the other genes associated with that trait out. The $P_{cond}$ reported in the text is the p-value of this gene-trait pair when adjusting for all other genes considered for conditioning for this trait, unless otherwise stated.

Table S6: Comparison of association results to independent data.

For each gene-trait pair, we list the original p-value, the GWAS trait(s) that we classified as most similar to a PheWAS trait, its best p-value in an independent dataset, the number of GWAS datasets that were used for this trait, and the rank of this gene within that dataset. For UK Biobank summary statistics, we have genome-wide data; for datasets with individual-level data, only 16p11.2 and 22q11.2 genes were calculated. See Table S2 for more information on datasets used.

Table S7: Traits over-represented in CNV carriers.

The four categories of CNV carrier – 16p11.2 duplication, 16p11.2 deletion, 22q11.2 duplication, 22q11.2 deletion – were tested separately. The results for all clinical traits tested are provided. The number of cases and controls for each trait is given, as well as whether the p-value meets either Bonferroni or FDR correction. Traits in bold were represented in over 5% of carriers.

Table S8: Top PheWAS associations of 16p11.2 and 22q11.2 genes.

The top 15 associated traits for each gene, regardless of p-value, are shown. These represent the top 1% of associations among all traits tested. Genes are listed in alphabetical order, with each trait's sample size and phecode (www.phewascatalog.org) noted.

Table S9: Enrichment of clinical categories among the top PheWAS associations.

The top 15 traits (codes) for each gene analyzed (n = 1470 gene-trait pairs) were divided into 17 clinical categories (observed counts column). The values in the expected counts column are calculated as 1470 * {the proportion of traits of that category tested}. For example, 159 out of 1531 codes tested were from the "circulatory system" category, so the expected counts for "circulatory system" are calculated as 1470*159/1531. The last column contains the p-value from a binomial test comparing whether the observed proportion of clinical categories is more extreme than expected.

Note S1: Supplemental Acknowledgements. Members of the Psychiatric Genomics Consortium who contributed to this work.

Figure S1

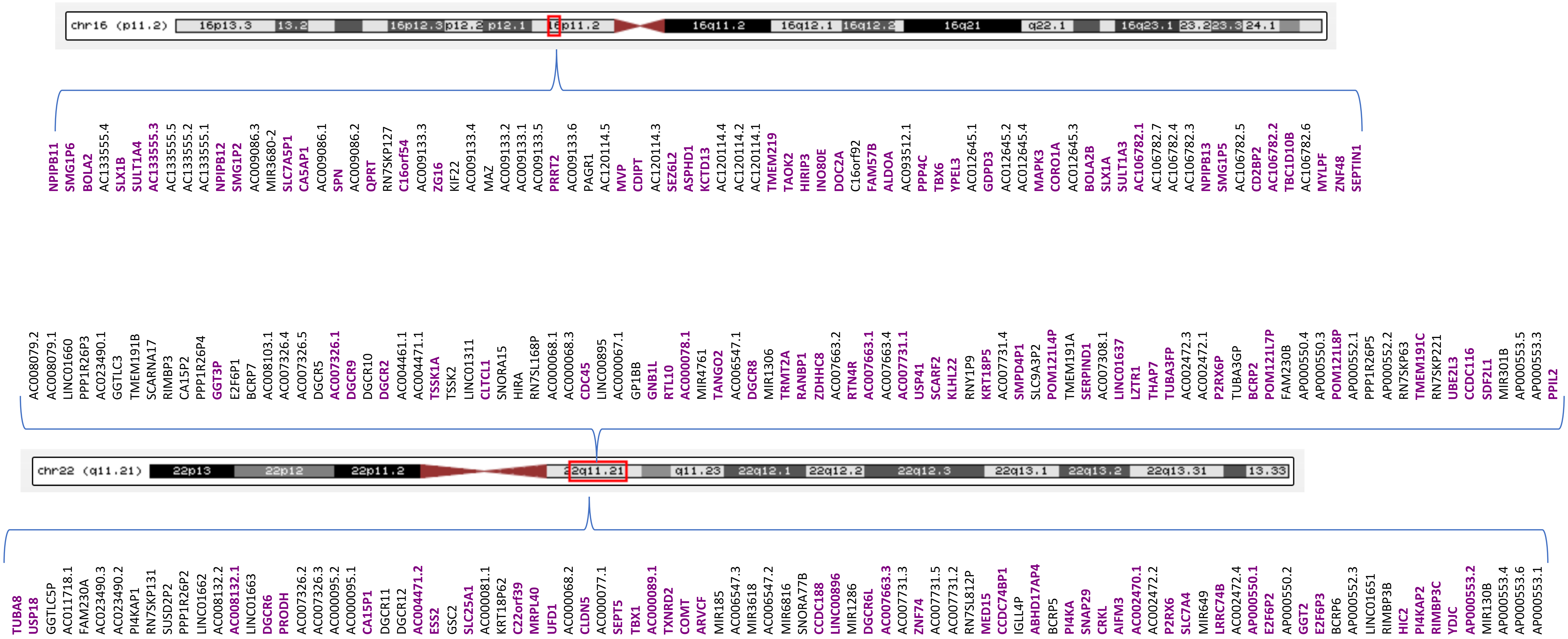
chr16 (p11.2)
16p13.3
13.2
16p12.3
p12.2
p12.1
16q11.2
16q12.1
16q12.2
16q21
q22.1
16q23.1
23.2
23.3
24.1
NPIPB11 SMG1P6 BOLA2 AC133555.4 SLX1B SULT1A4 AC133555.3 AC133555.5 AC133555.2 AC133555.1 NPIPB12 SMG1P2 AC009086.3 MIR3680-2 SLC7A5P1 CA5AP1 AC009086.1 SPN AC009086.2 QPRT RN7SKP127 C16orf54 AC009133.3 ZG16 KIF22 AC009133.4 MAZ AC009133.2 AC009133.1 AC009133.5 PRRT2 AC009133.6 PAGR1 AC120114.5 MVP CDIPT AC120114.3 SEZ6L2 ASPHD1 KCTD13 AC120114.4 AC120114.2 AC120114.1 TMEM219 TAOK2 HIRIP3 INO80E DOC2A C16orf92 FAM57B ALDOA AC093512.1 PPP4C TBX6 YPEL3 AC012645.1 GDPD3 AC012645.2 AC012645.4 MAPK3 CORO1A AC012645.3 BOLA2B SLX1A SULT1A3 AC106782.1 AC106782.7 AC106782.4 AC106782.3 NPIPB13 SMG1P5 AC106782.5 CD2BP2 AC106782.2 TBC1D10B AC106782.6 MYLPF ZNF48 SEPTIN1
AC008079.2 AC008079.1 LINC01660 PPP1R26P3 AC023490.1 GGTLC3 TMEM191B SCARNA17 RIMBP3 CA15P2 PPP1R26P4 GGT3P E2F6P1 BCRP7 AC008103.1 AC007326.4 AC007326.5 DGCR5 AC007326.1 DGCR9 DGCR10 DGCR2 AC004461.1 AC004471.1 TSSK1A TSSK2 LINC01311 CLTCL1 SNORA15 HIRA RN7SL168P AC000068.1 AC000068.3 CDC45 LINC00895 AC000067.1 GP1BB GNB1L RTL10 AC000078.1 MIR4761 TANGO2 AC006547.1 DGCR8 MIR1306 TRMT2A RANBP1 ZDHHC8 AC007663.2 RTN4R AC007663.1 AC007663.4 AC007731.1 USP41 SCARF2 KLHL22 RNY1P9 KRT18P5 AC007731.4 SMPD4P1 SLC9A3P2 POM121L4P TMEM191A SERPIND1 AC007308.1 LINC01637 LZTR1 THAP7 TUBA3FP AC002472.3 AC002472.1 P2RX6P TUBA3GP BCRP2 POM121L7P FAM230B AP000550.4 AP000550.3 POM121L8P AP000552.1 PPP1R26P5 AP000552.2 RN7SKP63 TMEM191C RN7SKP221 UBE2L3 CCDC116 SDF2L1 MIR301B AP000553.5 AP000553.3 PPIL2
chr22 (q11.21)
22p13
22p12
22p11.2
q11.23
22q12.1
22q12.2
22q12.3
22q13.1
22q13.2
22q13.31
13.33
TUBA8 USP18 GGTLC5P AC011718.1 FAM230A AC023490.3 AC023490.2 PI4KAP1 RN7SKP131 SUSD2P2 PPP1R26P2 LINC01662 AC008132.2 AC008132.1 LINC01663 DGCR6 PRODH AC007326.2 AC007326.3 AC000095.2 AC000095.1 CA15P1 DGCR11 DGCR12 AC004471.2 ESS2 GSC2 SLC25A1 AC000081.1 KRT18P62 C22orf39 MRPL40 UFD1 AC000068.2 CLDN5 AC000077.1 SEPT5 TBX1 AC000089.1 TXNRD2 COMT ARVCF MIR185 AC006547.3 MIR3618 AC006547.2 MIR6816 SNORA77B CCDC188 LINC00896 MIR1286 DGCR6L AC007663.3 AC007731.3 ZNF74 AC007731.5 AC007731.2 RN7SL812P MED15 CCDC74BP1 IGLL4P ABHD17AP4 BCRP5 PI4KA SNAP29 CRKL AIFM3 AC002470.1 AC002472.2 P2RX6 SLC7A4 MIR649 LRRC74B AC002472.4 AP000550.1 E2F6P2 AP000550.2 GGT2 E2F6P3 BCRP6 AP000552.3 LINC01651 RIMBP3B HIC2 PI4KAP2 RIMBP3C YDJC AP000553.2 MIR130B AP000553.4 AP000553.6 AP000553.1

Figure S2

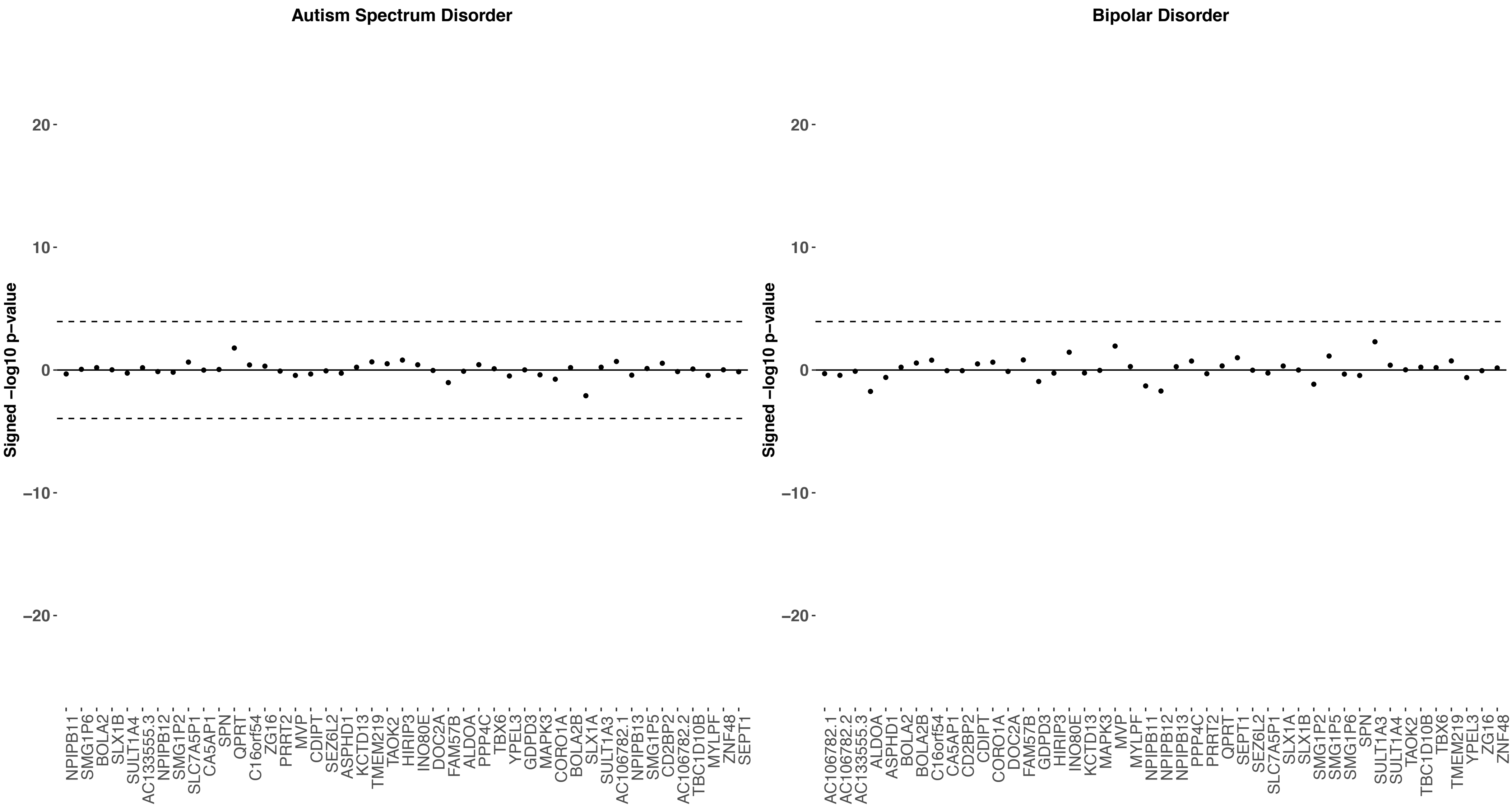

Autism Spectrum Disorder
Bipolar Disorder
Signed −log10 p−value
20
10
0
−10
−20
NPIPB11
SMG1P6
BOLA2
SLX1B
SULT1A4
AC133555.3
NPIPB12
SMG1P2
SLC7A5P1
CA5AP1
SPN
QPRT
C16orf54
ZG16
PRRT2
MVP
CDIPT
SEZ6L2
ASPHD1
KCTD13
TMEM219
TAOK2
HIRIP3
INO80E
DOC2A
FAM57B
ALDOA
PPP4C
TBX6
YPEL3
GDPD3
MAPK3
CORO1A
BOLA2B
SLX1A
SULT1A3
AC106782.1
NPIPB13
SMG1P5
CD2BP2
AC106782.2
TBC1D10B
MYLPF
ZNF48
SEPT1
AC106782.1
AC106782.2
AC133555.3
ALDOA
ASPHD1
BOLA2
BOLA2B
C16orf54
CA5AP1
CD2BP2
CDIPT
CORO1A
DOC2A
FAM57B
GDPD3
HIRIP3
INO80E
KCTD13
MAPK3
MVP
MYLPF
NPIPB11
NPIPB12
NPIPB13
PPP4C
PRRT2
QPRT
SEPT1
SEZ6L2
SLC7A5P1
SLX1A
SLX1B
SMG1P2
SMG1P5
SMG1P6
SPN
SULT1A3
SULT1A4
TAOK2
TBC1D10B
TBX6
TMEM219
YPEL3
ZG16
ZNF48

**Figure S3**

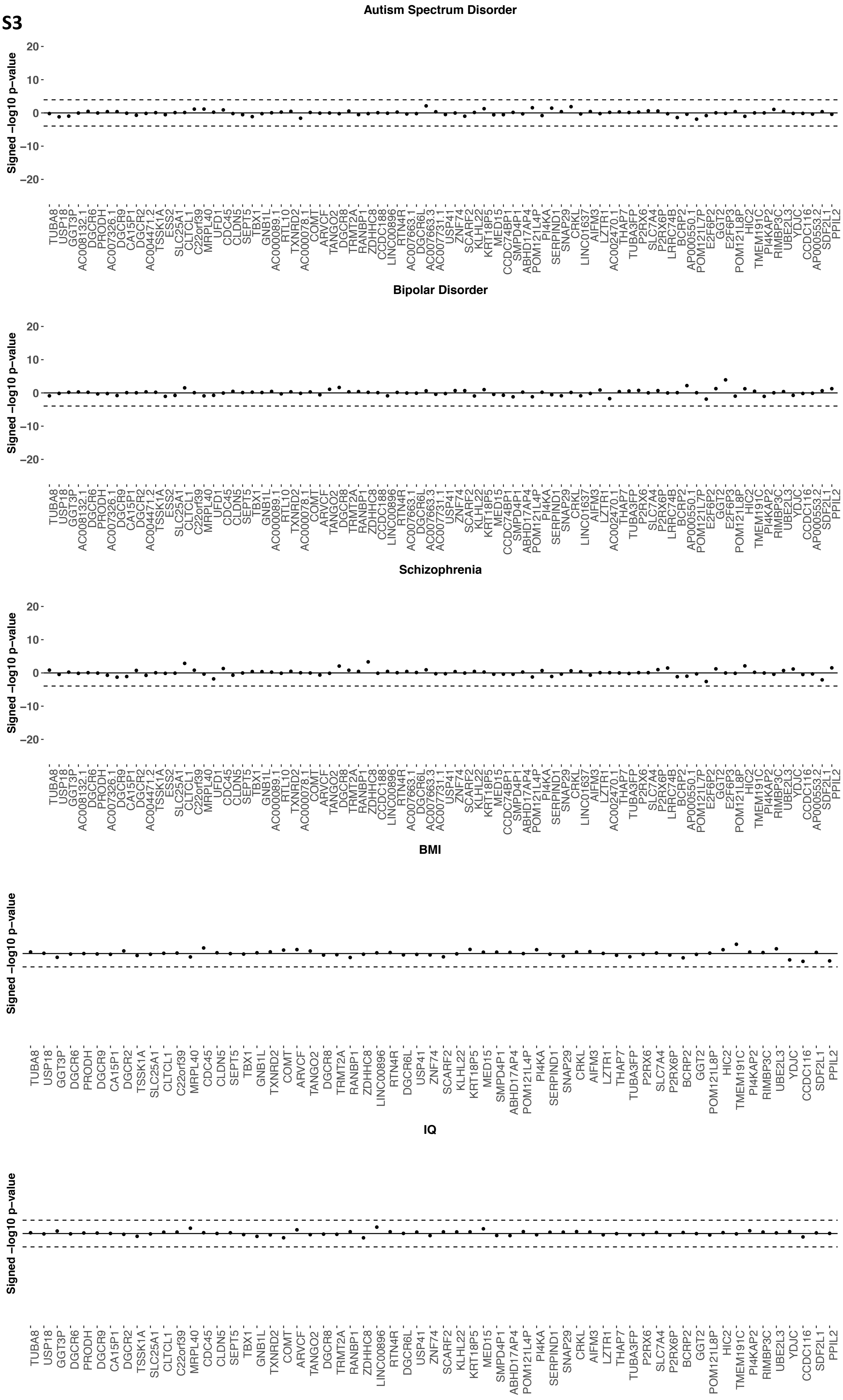
Autism Spectrum Disorder
Signed −log10 p−value
20
10
0
−10
−20
Bipolar Disorder
Signed −log10 p−value
Schizophrenia
Signed −log10 p−value
BMI
Signed −log10 p−value
IQ
Signed −log10 p−value

Figure S4

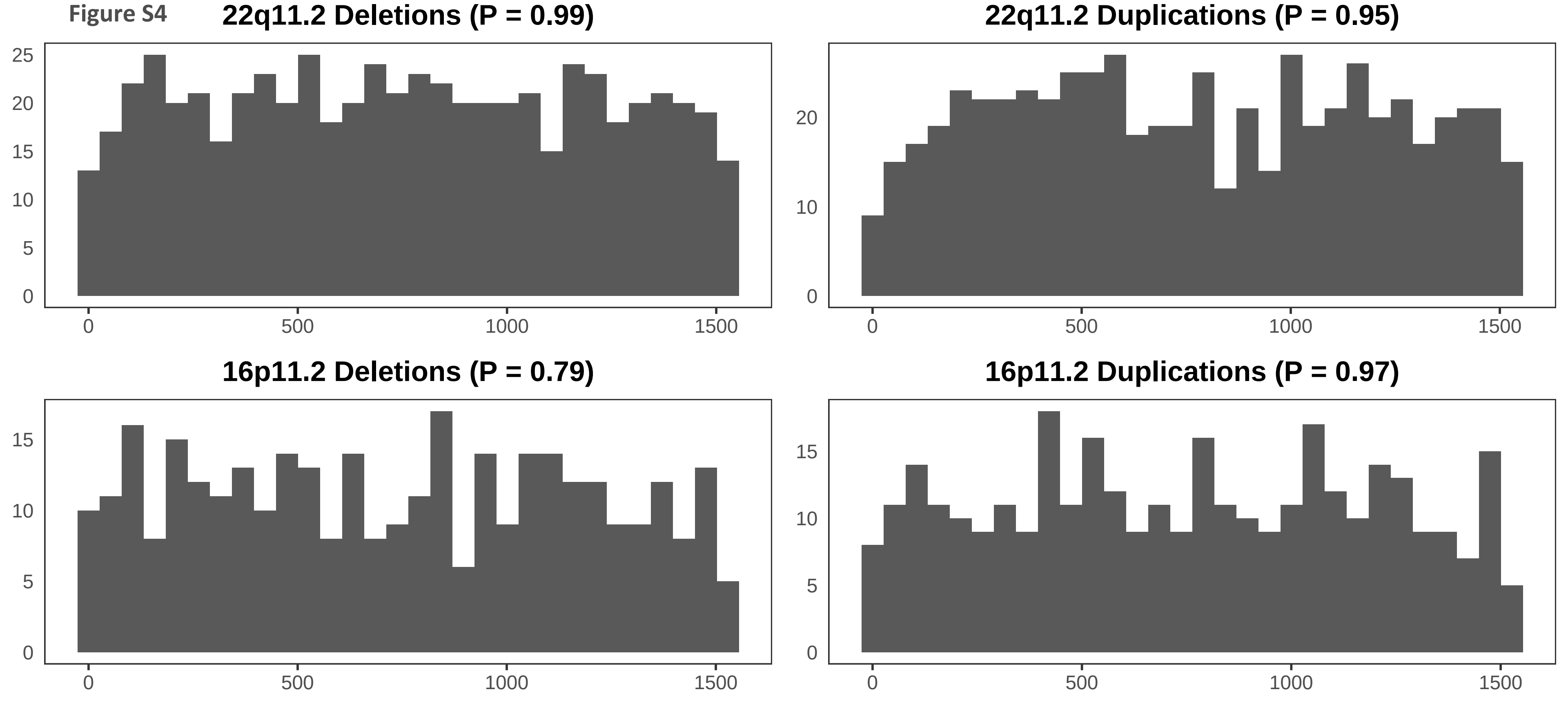

22q11.2 Deletions (P = 0.99)
22q11.2 Duplications (P = 0.95)
16p11.2 Deletions (P = 0.79)
16p11.2 Duplications (P = 0.97)
0
5
10
15
20
25
500
1000
1500

Note S1: Principal investigators of the Psychiatric Genomics Consortium

### Autism Working Group

Aravinda Chakravarti
Center for Human Genetics and Genomics, New York University School of Medicine, New York, New York, USA

Mark J, Daly
Stanley Center for Psychiatric Research, The Broad Institute of Harvard and M.I.T, Cambridge, MA, USA
Analytic and Translational Genetics Unit, Massachusetts General Hospital, Boston, USA
Institute for Molecular Medicine Finland, University of Helsinki, Helsinki, Finland

Daniel H, Geschwind
Department of Neurology and Center for Neurobehavioral Genetics, University of California Los Angeles, Los Angeles, CA

Joachim Franz Hallmayer
Department of Psychiatry and Behavioral Sciences, Stanford University School of Medicine, Stanford, CA

Aarno Palotie
Psychiatric and Neurodevelopmental Genetics Unit, Massachusetts General Hospital, Boston, MA, USA
Institute for Molecular Medicine Finland, FIMM, Helsinki, Finland
The Broad Institute of MIT and Harvard, Cambridge, MA, USA

Guy A. Rouleau
Department of Human Genetics, McGill University, Montreal, QC, Canada
Department of Neurology and Neurosurgery, Montreal Neurological Institute, McGill University, Montreal, QC, Canada

John E. Spiro
Simons Foundation, New York, New York

Joseph D. Buxbaum
Department of Human Genetics, Icahn School of Medicine at Mount Sinai, New York, NY, USA
Department of Psychiatry, Icahn School of Medicine at Mount Sinai, New York, NY, USA.
Friedman Brain Institute, Icahn School of Medicine at Mount Sinai, New York, NY, USA
Department of Neuroscience, Icahn School of Medicine at Mount Sinai, New York, NY, USA

Lauren A. Weiss
Department of Psychiatry, University of California San Francisco, San Francisco, CA, USA
Institute for Human Genetics, University of California San Francisco, San Francisco, CA, USA
Weill Institute for Neurosciences, University of California San Francisco, San Francisco, CA, USA

### Bipolar Working Group

Rolf Adolfsson
Department of Clinical Sciences, Psychiatry, Umeå University Medical Faculty, Umeå, SE

Ole A. Andreassen
Div Mental Health and Addiction, Oslo University Hospital, Oslo, NO
NORMENT, University of Oslo, Oslo, NO

Martin Alda, Gustavo Turecki, Guy A. Rouleau
Department of Psychiatry, Dalhousie University, Halifax, NS, CA;
National Institute of Mental Health, Klecany, CZ

Nicholas James Bass, Andrew McQuillin
Division of Psychiatry, University College London, London, GB

Joanna M. Biernacka, Mark Frye
Department of Health Sciences Research, Mayo Clinic, Rochester, MN, US
Department of Psychiatry & Psychology, Mayo Clinic, Rochester, MN, US

Douglas H. R. Blackwood
Division of Psychiatry, University of Edinburgh, Edinburgh, GB

Michael Boehnke, Laura J. Scott
Center for Statistical Genetics and Department of Biostatistics, University of Michigan, Ann Arbor, MI, US

Sven Cichon
Department of Biomedicine, University of Basel, Basel, Switzerland
Institute of Human Genetics, University of Bonn, School of Medicine & University Hospital Bonn, Bonn, Germany
Institute of Medical Genetics and Pathology, University Hospital Basel, Basel, Switzerland
Institute of Neuroscience and Medicine (INM-1), Research Centre Jülich, Jülich, Germany

Ashley L. Comes
Institute of Psychiatric Phenomics and Genomics (IPPG), University Hospital, LMU Munich, Munich, Germany
International Max Planck Research School for Translational Psychiatry (IMPRS-TP), Munich, Germany

Nicholas Craddock, Arianna Di Florio, Ian Jones
Medical Research Council Centre for Neuropsychiatric Genetics and Genomics, Division of Psychological Medicine and Clinical Neurosciences, Cardiff University, Cardiff, GB

Aiden Corvin
Neuropsychiatric Genetics Research Group, Dept of Psychiatry and Trinity Translational Medicine Institute, Trinity College Dublin, Dublin, IE

Franziska Degenhardt

Department of Child and Adolescent Psychiatry, Psychosomatics and Psychotherapy, University Hospital Essen, University of Duisburg-Essen, Duisburg, Germany
Institute of Human Genetics, University of Bonn, School of Medicine & University Hospital Bonn, Bonn, Germany

Andreas J. Forstner
Centre for Human Genetics, University of Marburg, Marburg, Germany
Institute of Human Genetics, University of Bonn, School of Medicine & University Hospital Bonn, Bonn, Germany

Janice M. Fullerton, Peter R. Schofield
Neuroscience Research Australia, Sydney, NSW, AU
School of Medical Sciences, University of New South Wales, Sydney, NSW, AU

Maria Grigoroiu-Serbanescu
Biometric Psychiatric Genetics Research Unit, Alexandru Obregia Clinical Psychiatric Hospital, Bucharest, Romania

José Guzman-Parra
Mental Health Department, University Regional Hospital, Biomedicine Institute (IBIMA), Málaga, Spain

Joanna Hauser
Department of Psychiatry, Laboratory of Psychiatric Genetics, Poznan University of Medical Sciences, Poznan, Poland

Lisa Jones
Department of Psychological Medicine, University of Worcester, Worcester, GB

John Kelsoe
Department of Psychiatry, University of California San Diego, La Jolla, CA, US

George Kirov
Medical Research Council Centre for Neuropsychiatric Genetics and Genomics, Division of Psychological Medicine and Clinical Neurosciences, Cardiff University, Cardiff, GB

Manolis Kogevinas
ISGlobal, Barcelona, Spain

Mikael Landén
Department of Medical Epidemiology and Biostatistics, Karolinska Institutet, Stockholm, SE
Institute of Neuroscience and Physiology, University of Gothenburg, Gothenburg, SE

Marion Leboyer
Faculté de Médecine, Université Paris Est, Créteil, FR
Department of Psychiatry and Addiction Medicine, Assistance Publique - Hôpitaux de Paris, Paris, FR
INSERM, Paris, FR

Jolanta Lissowska

Cancer Epidemiology and Prevention, M. Sklodowska-Curie National Research Institute of Oncology, Warsaw, Poland

Nicholas G. Martin
Genetics and Computational Biology, QIMR Berghofer Medical Research Institute, Brisbane, QLD, AU
School of Psychology, The University of Queensland, Brisbane, QLD, AU

Fermin Mayoral
Mental Health Department, University Regional Hospital, Biomedicine Institute (IBIMA), Málaga, Spain

Richard M. Myers
HudsonAlpha Institute for Biotechnology, Huntsville, AL, US

Philip B. Mitchell
Neuroscience Research Australia, Sydney, NSW, AU

Bertram Müller-Myhsok
Department of Translational Research in Psychiatry, Max Planck Institute of Psychiatry, Munich, Germany
Munich Cluster for Systems Neurology (SyNergy), Munich, Germany
University of Liverpool, Liverpool, United Kingdom

Markus M. Nöthen
Institute of Human Genetics, University of Bonn, School of Medicine & University Hospital Bonn, Bonn, Germany

Carlos N. Pato
Institute for Genomic Health, SUNY Downstate Medical Center College of Medicine, Brooklyn, NY, US
College of Medicine Institute for Genomic Health, SUNY Downstate Medical Center College of Medicine, Brooklyn, NY, US

Andreas Reif
Department of Psychiatry, Psychosomatic Medicine and Psychotherapy, University Hospital Frankfurt, Frankfurt am Main, Germany

Marcella Rietschel
Department of Genetic Epidemiology in Psychiatry, Central Institute of Mental Health, Medical Faculty Mannheim, Heidelberg University, Mannheim, DE

Sabrina Schaupp
Institute of Psychiatric Phenomics and Genomics (IPPG), University Hospital, LMU Munich, Munich, Germany

Lea Sirignano
Department of Genetic Epidemiology in Psychiatry, Central Institute of Mental Health, Medical Faculty Mannheim, Heidelberg University, Mannheim, Germany

Thomas G. Schulze

Institute of Psychiatric Phenomics and Genomics (IPPG), University Hospital, LMU Munich, Munich, Germany
Department of Psychiatry and Behavioral Sciences, Johns Hopkins University School of Medicine, Baltimore, MD, USA
Department of Genetic Epidemiology in Psychiatry, Central Institute of Mental Health, Medical Faculty Mannheim, Heidelberg University, Mannheim, Germany
Department of Psychiatry and Psychotherapy, University Medical Center Göttingen, Göttingen, Germany
Department of Psychiatry and Behaioral Sciences, SUNY Upstate Medical University, Syracuse, NY, USA

Jordan W. Smoller
Stanley Center for Psychiatric Research, Broad Institute, Cambridge, MA, US;
Department of Psychiatry, Massachusetts General Hospital, Boston, MA, US;
Psychiatric and Neurodevelopmental Genetics Unit (PNGU), Massachusetts General Hospital, Boston, MA, US

Fabian Streit
Department of Genetic Epidemiology in Psychiatry, Central Institute of Mental Health, Medical Faculty Mannheim, Heidelberg University, Mannheim, Germany

Patrick F. Sullivan
Department of Medical Epidemiology and Biostatistics, Karolinska Institutet, Stockholm, SE
Department of Genetics, University of North Carolina at Chapel Hill, Chapel Hill, NC, US
Department of Psychiatry, University of North Carolina at Chapel Hill, Chapel Hill, NC, US

Beata Świątkowska
Department of Environmental Epidemiology, Nofer Institute of Occupational Medicine, Lodz, Poland

### Schizophrenia Working Group

Rolf Adolfsson
Department of Clinical Sciences, Psychiatry, Umeå University Medical Faculty, Umeå, SE

Ole A. Andreassen
Div Mental Health and Addiction, Oslo University Hospital, Oslo, NO;
NORMENT, University of Oslo, Oslo, NO

Martin Begemann, Hannelore Ehrenreich, Agnes A. Steixner-Kuma
Clinical Neuroscience, Max Planck Institute of Experimental Medicine, Göttingen, Germany

Douglas H. R. Blackwood
Division of Psychiatry, University of Edinburgh, Edinburgh, GB

Anders D. Børglum
The Lundbeck Foundation Initiative for Integrative Psychiatric Research, iPSYCH, Denmark
Centre for Integrative Sequencing, iSEQ, Aarhus University, Aarhus, Denmark
Department of Biomedicine, Aarhus University, Aarhus, Denmark
Department P, Aarhus University Hospital, Risskov, Denmark.

Elvira Bramon
University College London, UK

Joseph D. Buxbaum
Department of Human Genetics, Icahn School of Medicine at Mount Sinai, New York, NY, USA
Department of Psychiatry, Icahn School of Medicine at Mount Sinai, New York, NY, USA.
Friedman Brain Institute, Icahn School of Medicine at Mount Sinai, New York, NY, USA
Department of Neuroscience, Icahn School of Medicine at Mount Sinai, New York, NY, USA

Aiden Corvin
Neuropsychiatric Genetics Research Group, Dept of Psychiatry and Trinity Translational Medicine Institute, Trinity College Dublin, Dublin, IE

Ariel Darvasi
Department of Genetics, The Hebrew University of Jerusalem, Jerusalem, Israel

Tõnu Esko
Medical and Population Genetics Program, Broad Institute of MIT and Harvard, Cambridge, MA, USA.
Division of Endocrinology and Center for Basic and Translational Obesity Research, Boston Children's Hospital, Boston, MA, USA
Department of Genetics, Harvard Medical School, Boston, MA, USA
Estonian Genome Center, University of Tartu, Tartu, Estonia

Pablo V. Gejman

Department of Psychiatry and Behavioral Neuroscience, University of Chicago, Chicago, IL, USA
Department of Psychiatry and Behavioral Sciences, NorthShore University HealthSystem, Evanston, IL, USA

Ina Giegling, Dan Rujescu
Department of Psychiatry, University of Halle, Halle, Germany

Christina M. Hultman
Department of Medical Epidemiology and Biostatistics, Karolinska Institutet, Stockholm,Sweden

Nakao Iwata
Department of Psychiatry, Fujita Health University School of Medicine, Toyoake, Aichi, Japan

Erik G. Jönsson
Department of Clinical Neuroscience, Karolinska Institutet, Stockholm, Sweden

George Kirov
Medical Research Council Centre for Neuropsychiatric Genetics and Genomics, Division of Psychological Medicine and Clinical Neurosciences, Cardiff University, Cardiff, GB

Todd Lencz
The Feinstein Institute for Medical Research, Manhasset, NY, USA
The Hofstra NS-LIJ School of Medicine, Hempstead, NY, USA
The Zucker Hillside Hospital, Glen Oaks, NY, USA

Douglas F. Levinson
Department of Psychiatry and Behavioral Sciences, Stanford University, Stanford, CA, USA.

Jianjun Liu
Human Genetics, Genome Institute of Singapore, A*STAR, Singapore
Saw Swee Hock School of Public Health, National University of Singapore, Singapore

Anil K. Malhotra
The Feinstein Institute for Medical Research, Manhasset, NY, USA
The Hofstra NS-LIJ School of Medicine, Hempstead, NY, USA
The Zucker Hillside Hospital, Glen Oaks, NY, USA

Bryan J .Mowry
Queensland Brain Institute, The University of Queensland, Brisbane, Queensland, Australia
Queensland Centre for Mental Health Research, University of Queensland, Brisbane, Queensland, Australia

Markus M. Nöthen
Institute of Human Genetics, University of Bonn, School of Medicine & University Hospital Bonn, Bonn, Germany

Michael C. O'Donovan, Michael J. Owen, James T.R. Walters

MRC Centre for Neuropsychiatric Genetics and Genomics, Institute of Psychological Medicine and Clinical Neurosciences, School of Medicine, Cardiff University, Cardiff, UK.
National Centre for Mental Health, Cardiff University, Cardiff, Wales

Roel A. Ophoff
Psychiatry, UMC Utrecht Brain Center Rudolf Magnus, Utrecht, NL
Human Genetics, University of California Los Angeles, Los Angeles, CA, US
Center for Neurobehavioral Genetics, University of California Los Angeles, Los Angeles, CA, US

Aarno Palotie
Psychiatric and Neurodevelopmental Genetics Unit, Massachusetts General Hospital, Boston, MA, USA
Institute for Molecular Medicine Finland, FIMM, Helsinki, Finland
The Broad Institute of MIT and Harvard, Cambridge, MA, USA

Carlos N. Pato
Institute for Genomic Health, SUNY Downstate Medical Center College of Medicine, Brooklyn, NY, US
College of Medicine Institute for Genomic Health, SUNY Downstate Medical Center College of Medicine, Brooklyn, NY, US

Tracey L. Petryshen
Department of Psychiatry, Harvard Medical School, Boston, MA, USA
The Broad Institute of MIT and Harvard, Cambridge, MA, USA
Center for Human Genetic Research and Department of Psychiatry, Massachusetts General Hospital, Boston, MA, USA

Marcella Rietschel
Department of Genetic Epidemiology in Psychiatry, Central Institute of Mental Health, Medical Faculty Mannheim, Heidelberg University, Mannheim, DE

Brien P. Riley
Virginia Institute for Psychiatric and Behavioral Genetics, Departments of Psychiatry and Human and Molecular Genetics, Virginia Commonwealth University, Richmond, VA, USA

Pak C. Sham
Centre for Genomic Sciences, State Ket Laboratory for Brain and Cognitive Sciences, and Department of Psychiatry, Li Ka Shing Faculty of Medicine, The University of Hong Kong, Hong Kong SAR, PR China

David St Clair
University of Aberdeen, Institute of Medical Sciences, Aberdeen, Scotland, UK.

Patrick F. Sullivan
Department of Medical Epidemiology and Biostatistics, Karolinska Institutet, Stockholm, SE
Department of Genetics, University of North Carolina at Chapel Hill, Chapel Hill, NC, US
Department of Psychiatry, University of North Carolina at Chapel Hill, Chapel Hill, NC, US

Daniel R. Weinberger

Lieber Institute for Brain Development, Baltimore, MD, USA
Departments of Psychiatry, Neurology, Neuroscience and Institute of Genetic Medicine, Johns Hopkins School of Medicine, Baltimore, MD, USA

Thomas Werge
The Lundbeck Foundation Initiative for Integrative Psychiatric Research, iPSYCH, Denmark.
Institute of Biological Psychiatry, MHC Sct. Hans, Mental Health Services Copenhagen, Denmark.
Department of Clinical Medicine, University of Copenhagen, Copenhagen, Denmark.
